# Biomineralized Surface-Enhanced Raman Scattering Nanotags Enable Machine Learning-Based Differentiation of Biomolecular Signatures

**DOI:** 10.64898/2026.05.23.726118

**Authors:** Md Hasnat Rashid, Moin Uddin Maruf, Troy Nations, Mahenour Megahed, Danielle E. Levitt, Balakrishna Koneru, Zeeshan Ahmad, Indrajit Srivastava

## Abstract

Surface-enhanced Raman scattering (SERS) nanotags provide highly sensitive platforms for in vitro diagnostics but often require complex, disease-specific customization that limits clinical translation. Biomineralization, in which biomolecules mediate inorganic material synthesis, offers a versatile yet underexplored strategy for generating functional SERS nanotags. Here, we demonstrate that biomolecule-directed biomineralization of gold nanoparticles (AuNPs) using amino acids and exosomes generates distinct nano-bio interfacial architectures that encode biomolecular identity into machine learning-resolvable SERS fingerprints through modulation of plasmonic coupling and Raman reporter organization. As a proof-of-concept system, amino acid-biomineralized AuNPs were synthesized using biomolecules with diverse physicochemical properties, including differences in size, polarity, and charge. The resulting nanotags were characterized using UV–Vis spectroscopy, SERS, fluorescence spectroscopy, dynamic light scattering (DLS), and transmission electron microscopy (TEM). Random forest and support vector machine (SVM) models successfully differentiated amino acid-dependent SERS signatures with near-perfect classification performance. Extending this approach to a biologically complex preclinical cancer model, exosome-biomineralized AuNP nanotags were generated using exosomes derived from clinically relevant pediatric patient-derived osteosarcoma and neuroblastoma tumors. Distinct exosome-dependent spectral fingerprints enabled SVM classification with 93.9% accuracy, while Shapley Additive exPlanations (SHAP) and t-distributed stochastic neighbor embedding (t-SNE) analyses identified diagnostically relevant spectral regions and visualized clustering between tumor classes. Collectively, this work establishes biomineralization as a strategy for transforming complex biomolecular and cellular information into computationally resolvable optical fingerprints, enabling scalable and label-free diagnostic classification of patient-derived biomolecular samples.

## INTRODUCTION

Early and accurate disease diagnosis remains fundamental to modern medicine. It enables timely intervention and improved patient outcomes.^1–4^ Surface-enhanced Raman scattering (SERS) has become a transformative analytical technique providing exceptional sensitivity, molecular specificity, and resistance to photobleaching.^5–7^ Although label-free SERS can amplify intrinsic Raman signals of analytes, it often fails to detect biomolecules with low Raman cross-sections, including certain proteins and lipids. To overcome this limitation, indirect SERS analysis employs “nanotags”, plasmonic nanoparticles functionalized with organic Raman reporters.^8, 9^ These have been extensively developed but nevertheless, the clinical translation of these nanotags is often limited. This is due to the need for complex, disease-specific customization and by challenges related to nanotag stability and signal variability in complex biological environments. ^5, 6, 10^

Biomineralization represents a promising yet underutilized strategy to address these challenges, offering a biologically inherent mechanism for the precision synthesis of functional nanomaterials.^11–14^ This process involves biological molecules or living systems mediating the nucleation, growth, and structural organization of inorganic materials under ambient conditions.^11,14^ In natural ecosystems, organisms employ tightly regulated biomineralization pathways to produce complex functional structures; classic examples include the formation of magnetosomes in magnetotactic bacteria for navigation ^15, 16^ and the sequestration of gold ions by species such as *Cupriavidus metallidurans* to mitigate heavy-metal toxicity.^17^ Beyond bacterial systems, this phenomenon spans various biomolecules. For instance, zygomycete fungi, such as *Rhizopus oryzae*, facilitate the reduction of gold(III) complexes into crystalline gold nanoparticles (AuNPs) via cell wall proteins ^18, 19^, while discrete proteins, such as lysozyme, can act as “nanoreactors” that template AuNP formation.^12, 20^ Unlike conventional “bottom-up” chemical syntheses that rely on harsh reducing agents, biomineralization proceeds under mild physiological conditions where the biomolecule acts simultaneously as the reducing agent, structural template, and stabilizing ligand.

This “green” synthesis route ensures that the resulting AuNPs retain a dense, natively derived bio-interface that preserves the intrinsic physicochemical properties, such as charge, polarity, and steric configuration, of the mediating biomolecules.^21^ This hybrid bio-nano architecture is critical for diagnostics, as demonstrated in recent studies on biomimetic SERS nanoparticles.^9^ Surface chemistry is the primary determinant of functional performance. In biomineralized systems, this unique biomolecular “shell” facilitates the differential attachment and orientation of Raman reporters on the AuNP surface. Because Raman signal intensity and peak ratios are highly sensitive to the proximity and orientation of the reporter relative to the plasmonic surface, the biomineralization agent serves as a spectroscopic determinant. Consequently, the SERS response provides a unique spectral fingerprint of the biological entity that mediated the nanoparticle growth. Importantly, these biomineralized spectral fingerprints likely arise from a combination of interconnected physicochemical effects rather than a single variable. Because the biomolecule directing AuNP formation remains associated with the nanoparticle surface, it establishes a distinct nano-bio interfacial environment that influences nanoparticle nucleation kinetics, surface charge distribution, local plasmonic coupling, and Raman reporter adsorption geometry.^8^ Variations in biomolecular polarity, steric organization, aromaticity, and metal-binding affinity can therefore modulate both electromagnetic hotspot formation and chemical interactions between the reporter and plasmonic surface.^22–24^ These coupled effects alter Raman peak intensities, relative band ratios, spectral variance, and enhancement efficiencies, ultimately generating high-dimensional SERS signatures characteristic of the biomineralizing species.

While cellular biomineralization has shown significant promise in therapeutic applications, such as the *in situ* generation of photothermal AuNPs within tumor environments ^25^, its potential as a diagnostic reporter of biological identity remains largely untapped. In this study, we present a scalable, label-free diagnostic framework that leverages biomineralization to convert complex biological information into distinguishable SERS signatures. For a schematic overview of our platform, see **Figure 1**. We demonstrate that the biomolecule-driven reduction of gold ions (Au^3+^ to Au^0^) imparts differentiable characteristics that allow for the high-fidelity discrimination of biomineralization agents through machine learning (ML) algorithms. As a proof of concept, we first established a library of SERS nanotags using amino acids encompassing a diverse range of physicochemical properties, including variations in size, polarity, and surface charge. These biomineralized platforms were rigorously characterized using UV-Vis absorbance spectroscopy, dynamic light scattering (DLS), zeta potential, transmission electron microscopy (TEM), and SERS spectroscopy. By employing ML classification algorithms such as random forests and support vector machine, we enabled the clear differentiation of these amino acid-derived spectroscopic signatures. Transitioning to a pre-clinical diagnostic model, we generated biomineralized SERS nanotags from exosomes derived from clinically relevant pediatric patient-derived osteosarcoma and neuroblastoma models. We show that their distinct spectral signatures form separate clusters in the t-distributed stochastic neighbor embedding (t-SNE) plots and the ML classification algorithms achieve an accuracy of 93.9% for this challenging task. Furthermore, we utilized interpretable ML (Shapley Additive exPlanation) analysis to identify specific wavenumbers with potential diagnostic and prognostic relevance. By leveraging the inherent chemical information of biomineralizing agents, this platform provides a label-free framework for the molecular analysis of clinical samples, integrating green nanotechnology with advanced SERS-based diagnostics.

**Figure 1.**
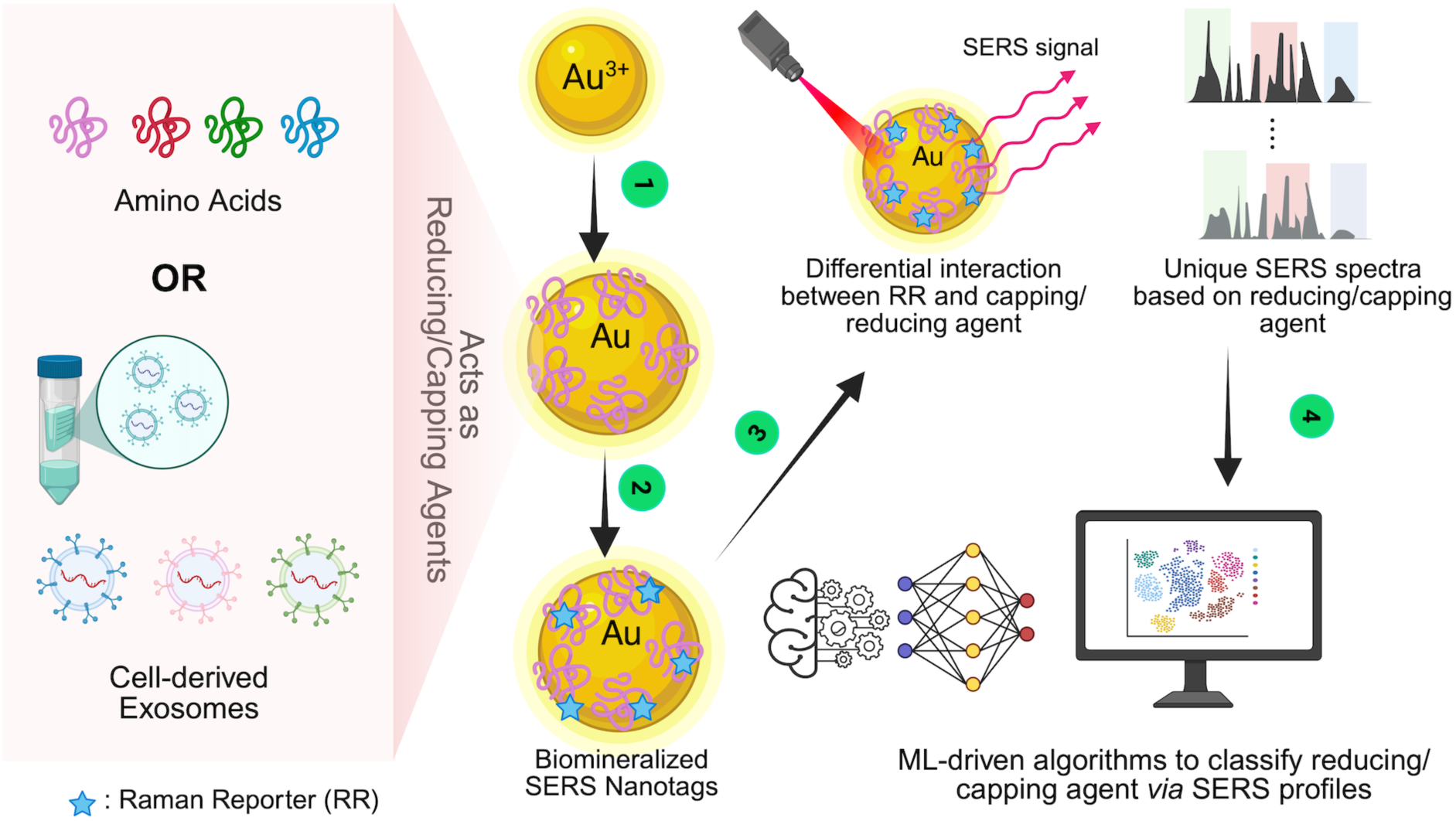
Schematic illustration of the biomineralization and SERS-AI platform for biomolecular sensing. **(1)** AuNPs are synthesized *via* biomimetic reduction using various capping agents, including -amino acids, proteins, and tumor cell-derived exosomes. **(2)** Functionalization with a Raman reporter molecule generates biomineralized SERS nanotags. **(3)** Upon laser excitation, the unique surface chemistry of each capping agent modulates the molecular orientation of the reporter, yielding distinct SERS “fingerprints.” **(4)** High-dimensional spectral data are processed via AI/ML models to achieve high-accuracy classification, enabling the discrete separation of all seven amino acid-stabilized AuNPs and the lineage-specific differentiation between neuroblastoma and osteosarcoma-derived exosomes.

## EXPERIMENTAL SECTION

### Reagents and Materials

dl-phenylalanine (C_9_H_11_NO_2_, 99%), l-proline (C_5_H_9_NO_2_, 99%), l-valine (C_5_H_11_NO_2_, 99%), l-serine (C_3_H_7_NO_3_, 99%), l-glutamic acid (C_5_H_9_NO_4_, 99%), l-Glycine (C_2_H_5_NO_2_, 98.5% p.a.). L-leucine (C_6_H_13_NO_2_, 99%) and gold (III) chloride tri-hydrate (HAuCl_4_·3H_2_O, 90% Au) were purchased from Sigma-Aldrich. Sodium hydroxide (NaOH, 99%), phosphate-buffered saline (PBS; pH 7.4), and Invitrogen total exosome isolation reagent kit (from cell culture media) were purchased from Thermo Fisher Scientific. Deionized (DI) water was used throughout the experiments. All glassware was treated before the reactions with aqua regia and subsequently rinsed several times with DI water before use.

### Synthesis of *α*-Amino Acid biomineralized Gold Nanoparticles (AA-AuNP)

AuNP solutions were prepared using a method described by Turkevich and were successfully adjusted for AuNP synthesis using different amino acids.^26^ In a typical procedure, 1.5 mL of 5mM HAuCl_4_ aqueous solution was added to 27 mL of water and was heated to 100 °C on a hot plate, stirring at 450 rpm. Then, 1.5 mL of α-amino acid solution was quickly added to the mixture. After that, 50-200 µL of 0.1M freshly produced NaOH was added to the mixture, and the reaction was performed until the solution acquired a distinctive color (red, pink, or purple) that was stable for at least 5 minutes.

### Cell Culture Protocol

Neuroblastoma patient-derived cell lines SK-N-BE(2) and COG-N-519 were obtained from the ALSF/Children’s Oncology Group (COG) Childhood Cancer Repository (www.CCcells.org). The osteosarcoma cell line U2OS was obtained from the American Type Culture Collection (ATCC), and the OS526 cell line was provided by the laboratory of Alejandro Sweet-Cordero at University of California, San Francisco. All patient-derived cell lines were established and maintained under physiologic hypoxic culture conditions consisting of 5% oxygen and 5% CO_2_ and were cultured in antibiotic-free Iscove’s Modified Dulbecco’s Medium supplemented with 20% fetal bovine serum (FBS), 1× insulin transferrin selenium (ITS), and 4 mM L-glutamine. Human cell line identity was validated using the GenePrint 10 System (Promega, Madison, WI). For exosome isolation experiments, conditioned supernatant media was collected from tissue culture flasks 96 hours after cell seeding. Prior to collection, cells were maintained under standard growth conditions to allow accumulation of extracellular vesicles and secreted factors within the culture media, while minimizing cellular over confluency and excessive cell death.

### Exosome Isolation from Cell Culture Supernatant using ultra-centrifugation approach

Exosomes from COG-N-519 and SK-N-BE(2) cell culture supernatant were isolated by differential centrifugation. ^27, 28^ 20 mL of culture supernatant was first centrifuged at 2000 × g for 10 minutes at room temperature to pellet out cell debris. The pellet was discarded, and the resulting supernatant was centrifuged at 15,500 × g for 30 minutes at 4 °C to pellet out larger particles. The pellet was again discarded, and the supernatant was filtered through a 0.22 µm PVDF syringe filter into ultracentrifuge (UC) tubes and ultracentrifuged at 100,000 × g for 90 min at 4 °C (Himac CS150Fnx, Eppendorf, Hamburg, Germany). Following the first UC step, all but 1 mL supernatant was removed, and the pellets were resuspended in remaining supernatant by pipetting. After transferring resuspended pellets to a 1.5 mL polycarbonate microcentrifuge tube, samples were ultracentrifuged at 100,000 × g for 70 min at 4 °C. The supernatant was removed, and the pellet was resuspended in 100 µL of 0.02 µm-filtered PBS for downstream applications,

### Exosome Isolation from Cell Culture Supernatant using The Invitrogen total Exosome Isolation Kit

Exosomes from OS-526 and U20S cell culture supernatants were isolated using a Invitrogen total exosome isolation (for cell culture media) kit following the manufacturer’s recommended protocol. First, 20 mL of culture supernatant was centrifuged at 2000 × g for 30 min at 4 °C using Sorvall ST4 Plus Centrifuge to remove cell debris. Then, the pellet was discarded, and the supernatant was transferred into a new tube where 0.5 volumes of total exosome isolation reagent were added. The mixture was mixed thoroughly and incubated at 2-8 °C overnight. Then, the mixture was centrifuged at 10,000 × g for 1 hour at 4°C. Thereafter, the supernatant was discarded, and the pellets were resuspended in 1mL of sterile PBS for downstream applications.

### Synthesis of Exosome biomineralized Gold Nanoparticles (Exo-AuNP

Exo-AuNPs were synthesized following a modified protocol from Lee et al.^29^ Initially, isolated exosomes were thawed overnight at 4 °C before they were used in the experiment. Briefly, 2 mL of 1.25 mM HAuCl_4_ was mixed with 2 mL isolated exosome solution diluted in PBS 1X and stirred at room temperature for 5 minutes to allow for ion coordination. Then the pH of the solution was adjusted to 11.5 using 0.1M NaOH to initiate the reduction process. The mixture was then incubated at 37°C in a shaking incubator (200 rpm) for 24 hours. Successful synthesis was monitored via a characteristic color transition from light yellow to colorless, and finally to a stable pink. After incubation, the resulting Exo-AuNPs were stored 4°C for subsequent characterization.

### Physiochemical Characterization of SERS Nanotags

UV-visible absorption spectra of the synthesized AA-AuNPs or Exo-AuNPs was measured by GENESYS™ 30 Visible Spectrophotometer (Thermo Fisher) at 350-800 nm wavelength without any dilutions. Hydrodynamic diameter and Zeta potential were measured by DLS Litesizer 700 (Anton Parr) with a sample ratio of 50 *μ*L:950 *μ*L with DI water utilizing prior established protocols.^30, 31^ ^32^ Transmission electron microscopy (TEM) imaging was performed following a previously reported negative-staining protocol.^33, 34^ Briefly, a 25 μL drop of the liquid sample was deposited onto a glow-discharged, carbon-coated copper Ted Pella 01801 TEM grid and incubated for 1 minute. After incubation, the excess liquid was blotted off using filter paper, and the grid was dried in a vacuum desiccator for four hours. The sample-coated grid was immediately counterstained with 1% uranyl acetate (UA) for 30 seconds. Excess UA was then blotted off, and the grid was dried in a vacuum desiccator for four hours before imaging. All samples were imaged using a Hitachi S-7650 transmission electron microscope operated at 100 kV.

### BCA Assay of Exosomes and Exo-AuNPs

The protein concentration in the exosomes and Exo-AuNPs was quantified using a bicinchoninic acid assay (BCA) kit (Pierce™ BCA Protein Assay Kit, Thermo Fisher Scientific) according to the manufacturer’s protocol. Briefly, an 11-point bovine serum albumin (BSA) standard curve (0–2000 µg/ml) was prepared by serial dilution using 1x PBS. Then, 25 µl of each standard and samples were added per well in triplicates. A working reagent was prepared using BCA reagent A and B mixed at a ratio of 50:1 and 200 µl of the working reagent was added to each well. The plate was then incubated at 37°C for 30 min, after which the absorbance was read using BioTek Epoch 2 Microplate Spectrophotometer (Agilent) at 562 nm. The protein concentrations of the samples were interpolated from the standard curve.

### SDS PAGE of Exosomes and Exo-AuNPs

The size distribution of proteins in the exosomes and Exo-AuNPs samples were visualized using sodium dodecyl sulfate–polyacrylamide gel electrophoresis (SDS–PAGE). 30 μl of the samples were dissolved in 4x Laemmli buffer (Bio-Rad) containing 10% β-mercaptoethanol (Bio-Rad) and heated at 95°C for 8 minutes. The samples were centrifuged for 20 seconds at 3000 rpm before being loaded and resolved onto 10% Mini-PROTEAN TGX precast gels (Bio-Rad) for 1 hour at 200 V. Following electrophoresis, the gels were stained with Coomassie Brilliant Blue G-250 (Bio-Rad) for 1 hour followed by washing in distilled water for 1 h. The distribution of proteins was visualized using ChemiDoc Go Imaging System (Bio-Rad).

### SERS Spectra Acquisition

For SERS label incorporation, 2 mL of AA-AuNP or Exo-AuNP solution was transferred to a glass vial, and as a Raman reporter, IR-780 iodide dye was added at volumes determined through prior concentration optimization: 8 µL for AA-AuNPs and 4 µL for Exo-AuNPs, corresponding to the concentrations yielding maximum Raman signal intensity without spectral saturation or aggregation-induced peak broadening. The mixture was vortexed for 2 minutes to ensure uniform dye adsorption onto the nanoparticle surface, then allowed to equilibrate for 5 minutes at room temperature before spectral acquisition. SERS measurements were performed using a Cora 5000 Raman Spectrometer (Anton Paar) equipped with a 785 nm excitation laser. Instrument parameters were independently optimized for each nanotag formulation: AA-AuNP spectra were acquired at 200 mW laser power with a 200 ms exposure time, while Exo-AuNP spectra were collected at 50 mW with a 100 ms exposure. The resulting spectra were baseline-corrected and preprocessed prior to machine learning classification.

### Evaluation of SERS signal stability

To determine the optimal IR-780 dye loading concentration, 2mL aliquots of the AA-AuNP suspension were transferred to a glass vial, and IR- 780 was added in incremental volumes ranging from 0 to 20 µL. Each sample was then vortexed for two minutes and allowed to equilibrate for 5 minutes at room temperature. After that, their SERS spectra were collected using the previously mentioned parameters. To evaluate their temporal signal stability, mixtures of AA-AuNP with IR-780 dye were stored under ambient conditions, and their spectra were measured on days 1, 2, and 3 post-preparations. This spectra was then processed and plotted to evaluate their signal stability over a three-day period.

### Dataset Splits and Classifier Configuration

For the amino acid analysis, each of the 7 amino acids contributed 80 samples. For the ratiometric analysis, 20 spectra were collected for each proline:valine mixture. For the exosome analysis, the dataset comprised 115, 103, 113, and 109 spectra for Exo-AuNP@COG-N-519, @OS-526, @SK-N-BE(2), and @U2OS, respectively. All of the datasets were partitioned into 70% training, 15% validation, and 15% test sets via random sampling. Because the splits were drawn randomly without stratification, the per-class counts in the test set are not identical across amino acids which is reflected in the diagonal entries of the confusion matrices (Fig. 3a, Fig. 4a, and Fig. 5d,e). An additional 20 samples for 4 amino acids were held out as a blind test set. The model’s performance is evaluated on this blind set separately. We used random forest and support vector machine algorithms to classify the spectra. Both classifiers were used as implemented in scikit-learn with fixed random seeds for reproducibility.^35^

### ML Classification Algorithms

#### Random Forest

Random Forest (RF) is an ensemble learning method that constructs a multitude of decision trees during training and outputs the class receiving the most votes across all trees. Originally introduced by Breiman ^36^, RF builds each tree on a bootstrapped subset of the training data and considers only a random subset of features at each split, reducing overfitting and improving generalization compared to single decision trees. Its intrinsic feature importance metric, derived from the mean decrease in impurity across all trees, makes it particularly well-suited for high-dimensional spectroscopic data where identifying discriminative wavenumbers is as important as classification accuracy itself. RF has been demonstrated to be highly compatible with SERS data analysis, where its ability to identify important spectral variables and analyze inter-variable relationships enables accurate and interpretable classification of complex biological samples.^37^ The RF classifier we used was configured with n_estimators = 100 and random_state = 42.

#### Support Vector Machine

Support Vector Machine (SVM) is a kernel-based supervised classifier that identifies the optimal hyperplane maximizing the margin between classes in a transformed feature space, making it particularly effective for datasets where classes are not linearly separable in their original dimensionality.^38^ The use of the radial basis function (RBF) kernel allows SVM to capture nonlinear decision boundaries, which is advantageous when SERS spectra of biologically similar samples differ in subtle intensity patterns rather than discrete peak presence or absence. SVM has demonstrated strong performance in biomedical spectral classification tasks, particularly for exosome and cell-line discrimination. Liu et al. employed a PCA-SVM framework applied to SERS spectra of exosomes derived from six distinct cell lines - HepG2, HeLa, 143B, LO-2, BMSC, and H8, achieving a classification accuracy of 94.4% using only a small number of spectra per class.^39^ Similarly, Liu et al. applied a Bayesian-optimized SVM model to SERS spectra of lung cancer cell-derived extracellular vesicles, achieving a cross-validation loss of only 3.7% and an independent test set accuracy of 98.7%, with the approach further validated on plasma-derived exosomes from both mouse models and human patients.^40^ The Support Vector Machine used an RBF kernel with regularization parameter C = 1, gamma = ‘scale’, and random_state = 42. Collectively, ML techniques such as SVM and RF are generally more interpretable and require less computational power than deep learning approaches, making them particularly suitable for small to moderate spectral datasets and structured SERS data in exosome-based diagnostic workflows.

### ML interpretability algorithms

#### t-distributed stochastic neighbor embedding (t-SNE)

To visualize the high-dimensional spectral data in two dimensions, we applied t-distributed Stochastic Neighbor Embedding (t-SNE), a nonlinear dimensionality reduction technique introduced by van der Maaten and Hinton that preserves local neighborhood structure by modeling pairwise similarities as conditional probabilities and minimizing the Kullback–Leibler divergence between the high- and low-dimensional distributions.^41^ t-SNE is particularly well-suited for revealing cluster structure in SERS datasets, where class separability is often nonlinear. We used the scikit-learn implementation with n_components = 2, perplexity = 30, and random_state = 42 to ensure reproducibility, where the perplexity parameter balances attention between local and global aspects of the data. ^35^

#### SHAP

To interpret the contribution of individual wavenumbers to each classifier’s predictions, we applied SHapley Additive exPlanations (SHAP), a unified, game-theoretic framework introduced by Lundberg and Lee that assigns each feature an importance value reflecting its marginal contribution to a given prediction.^42^ For the Random Forest classifier, we used shap.TreeExplainer, which exploits the tree structure to compute exact Shapley values in polynomial time.^43^ To make the kernel-based estimation computationally tractable, the training set was summarized into a 30-cluster background distribution using shap.kmeans(X_train_sc, 30), and SHAP values were computed on a subset of 24 test samples with nsamples = 100 Monte Carlo perturbations per explanation.

## RESULTS AND DISCUSSION

### Biomineralization of AuNPs *via* Amino Acids Side-Chain Diversity

AuNPs were synthesized *via* a biomimetic reduction process termed as biomineralization. In this approach, specific biomolecules, including amino acids proteins, or exosomes mediate the reduction of gold ions (Au^3+^ to Au^0^), facilitating the controlled nucleation and growth of nanoparticles with tunable physicochemical properties. To investigate the fundamental role of molecular structure in the process, we first employed seven distinct alpha amino acids as dual reducing and stabilizing agents: L-glutamic acid (GLU), L-glycine (GLY), L-leucine (LEU), DL-phenylalanine (PHE), L-proline (PRO), L-serine (SER), and L-valine (VAL) (**Figure 2a, b**). While these amino acids share a conserved structural backbone, comprising an amino group (–NH_2_), a carboxyl group (–COOH), and a central α-carbon, their divergent side chain (R-groups) fundamentally alter their chemical behavior. These R-groups dictate the polarity, hydrophobicity, and metal-binding affinity of the local biochemical environment during synthesis.^44–47^ For instance, the polar side chains of L-glutamic acid and L-serine promote hydrogen bonding and coordination with gold ions, whereas the nonpolar residues of L-leucine and L-valine provide hydrophobic stabilization. Furthermore, the unique cyclic structure of L-proline imposes steric constraints on the particle surface, while bulky aromatic ring of DL-phenylalanine influences the electronic density and orientation of surface-bound molecules. These specific chemical interactions result in the formation of distinct AuNPs, each characterized by unique morphologies, size distributions, and spectroscopic behaviors (**Figure 1**). Such variability is central to the performance of the resulting SERS nanotags. By leveraging the side-chain diversity, we developed a tunable platform where distinct spectroscopic signatures enable high-accuracy classification through ML-driven analysis, opening up a potential for robust strategy for biomolecule-specific sensing applications.

**Figure 2.**
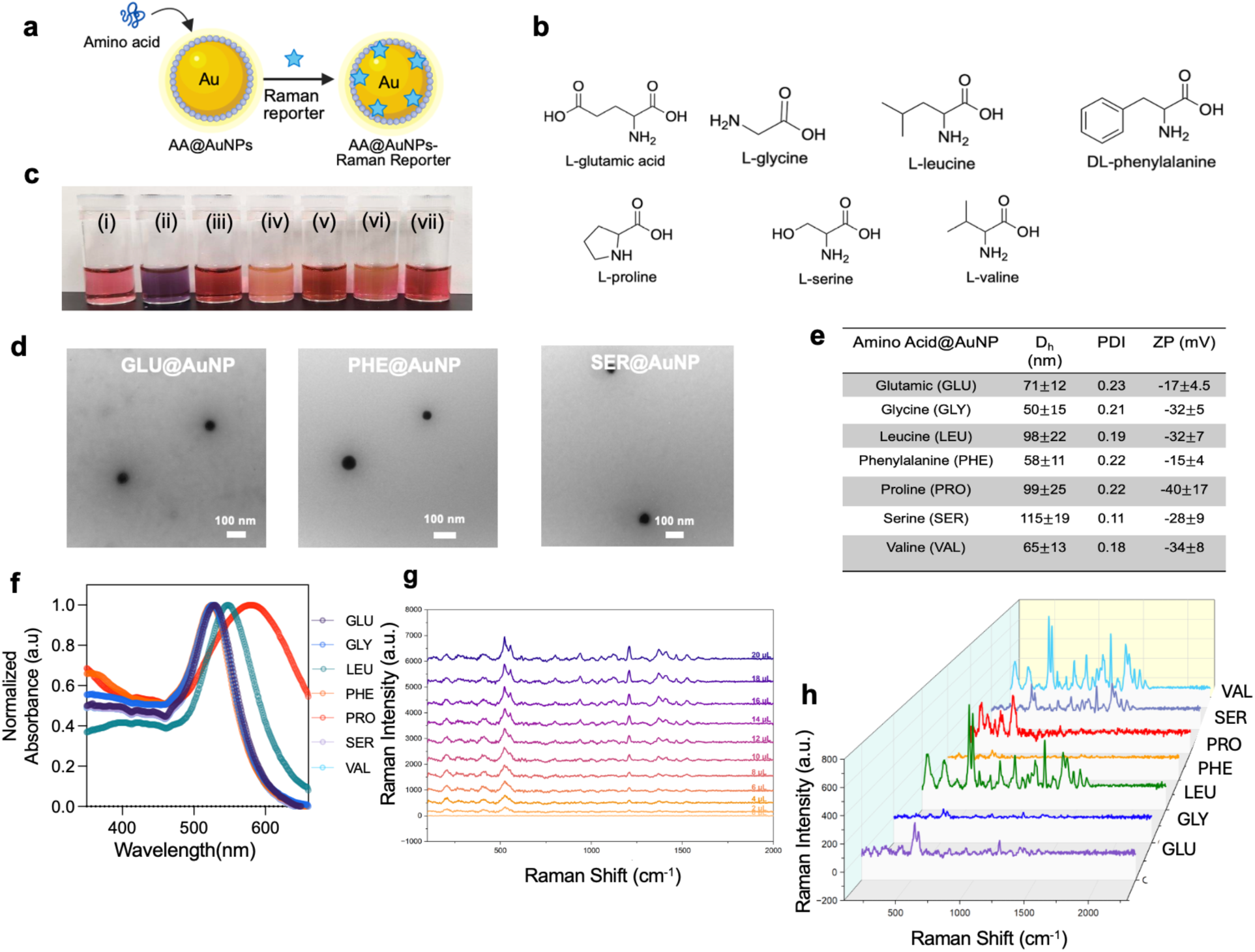
Synthesis and characterization of amino acid-biomineralized gold nanoparticles (AA@AuNPs). **(a)** Schematic representation of AuNP biomineralization using a-amino acids followed by the addition of a Raman reporter to create SERS nanotags. **(b)** Chemical structures of the seven a-amino acids used in this study. **(c)** Digital photographs of the biomineralized AuNP solutions: (i) Glutamic Acid, (ii) Proline, (iii) Glycine, (iv) Serine, (v) Phenylalanine, (vi) Leucine, and (vii) Valine. **(d)** Representative Transmission Electron Microscopy (TEM) images of selected AA@AuNPs (scale bars=100 nm). **(e)** Physicochemical characterization including hydrodynamic diameter (D_h_), polydispersity index (PDI), and zeta potential (ZP). **(f)** UV-Vis absorbance spectra displaying the localized surface plasmon resonance (LSPR) peaks of AA@AuNPs. **(g)** SERS intensity of representative Glu@AuNP as a function of Raman reporter concentration (2–20 *μ*L). **(h)** Distinct SERS fingerprint spectra were collected for the seven different AA-AuNP nanotags.

The biomineralization process produced AA-AuNPs with distinct colorimetric profiles (**Figure 2c**), reflecting variations in their particle size and surface charge (**Figure 2e).** Dynamic light scattering confirmed the hydrodynamic diameters (D_h_) ranged from 50 nm to 115 nm, with Ser-AuNPs exhibiting the largest diameter. A plausible explanation for these variations could be attributed to the diverse side-chain polarities and steric profiles of the amino acids, which dictate nucleation and growth kinetics. Zeta potential (ζ) measurements ranged from -40 mV to -15 mV for all samples, indicating robust electrostatic stabilization.^47^ Some AA-AuNPs exhibited less negative zeta potential, such as Glu-AuNPs and Phe-AuNPs, likely due to specific side-chain coordination modes that offer less effective charge screening. Despite these differences, polydispersity indexes (PDI) remained between 0.10 and 0.23, confirming the formation of monodisperse systems essential for reproducible SERS performance. Representative transmission electron microscopy (TEM) (**Figure 2d, S1-7**) corroborated these findings, revealing well-dispersed, spherical morphologies. As expected, values from DLS were consistently larger than the core diameters observed in TEM as DLS accounts for the hydration shell and surface-bound ligands in solution while TEM measures the dehydrated metallic NPs in a dry state. UV-Vis absorption spectra (**Figure 2f**) further showed characteristics surface-plasmon resonance (SPR) peaks within the 520 – 600 nm range. The narrow SPR peaks suggestion uniform dispersion, with a slight broadening for Pro-AuNPs potentially resulting from the constrained cyclic structure of proline affecting stabilization.

To generate SERS-active nanotags, the AuNPs were functionalized with IR-780 iodide, a near-infrared resonant Raman reporter.^6^ ^9, 48^ The extended *π*conjugated system and positive charge of the IR-780 molecules enable robust absorption onto the AuNPs via cooperative electrostatic interactions and *π*- *π* stacking. However, the unique surface chemistry of each AA-AuNP, defined by the specific amino acid used during biomineralization, acts as a molecular template that governs these interactions. Variations in side-chain polarity, aromacity, and steric configuration create distinct chemical environments ^45, 46^ that modulate the final molecular orientation and packing density of the IR-780 dye within the plasmonic “hot spots”. Consequently, while the reporter remains identical across all samples, its specific interaction with the divergent amino acid shell results in distinctly different SERS fingerprints (**Figure 2h**). This variability confirms that the biomineralized outer layer does not merely stabilize the particle, but actively dictates the optical output, providing the high-dimensional data platform necessary for ML-based classifications. As demonstrated by the concentration titration in **Figure 2g**, the SERS signal intensity increases proportionally with dye concentration, confirming that the observed spectral diversity is a result of these specific surface interactions rather than inconsistent dye loading. To optimize reporter loading, SERS signal intensity was evaluated as a function of increasing IR-780 concentration (0–20 µL) for each AA-AuNP nanotag formulation **(Figures S8–S13)**. The temporal stability of the SERS signal was further evaluated by collecting spectra over three days following reporter incorporation. Consistent spectral profiles was observed across all time points, confirming reliable signal reproducibility.

The biomineralized AA-AuNP nanotags exhibited substantial differences in both SERS enhancement intensity and spectral morphology despite incorporation of an identical IR-780 Raman reporter across all formulations. The relative enhancement trend was approximately VAL ≈ LEU > SER > GLU > PRO > GLY ≈ PHE, indicating that the amino acid-mediated nano-bio interface strongly modulates reporter adsorption and plasmonic coupling. VAL-AuNPs and LEU- AuNPs produced the strongest enhancement, potentially due to hydrophobic interactions that promote closer association of the hydrophobic IR-780 reporter with the Au surface and increase accessibility to plasmonic hotspots. SER-AuNPs also exhibited strong enhancement, suggesting that hydroxyl-mediated hydrogen bonding may support favorable reporter organization near the nanoparticle interface. In contrast, GLU-AuNPs generated only moderate enhancement despite the negatively charged glutamate surface potentially attracting the positively charged IR-780 reporter electrostatically, indicating that strong surface association alone does not necessarily maximize SERS activity. Excessive or diffuse electrostatic adsorption may orient the reporter outside regions of maximal electromagnetic enhancement or reduce effective hotspot coupling. PRO-AuNPs displayed intermediate spectral intensity, potentially due to steric constraints imposed by the cyclic proline structure. GLY-AuNPs and PHE-AuNPs generated the weakest spectra, likely through distinct mechanisms. Glycine lacks a substantial side chain capable of directing organized reporter packing, whereas the bulky aromatic phenylalanine side chain may hinder optimal reporter orientation relative to plasmonic hotspots. Collectively, these findings demonstrate that biomineralization actively governs nanoscale reporter organization and electromagnetic coupling at the nano-bio interface, thereby generating distinct amino acid-dependent SERS fingerprints.

### High Accuracy Classification of AA-AuNPs and Single-Blind Validation *via* Machine Learning

To determine whether the amino acid-dependent SERS fingerprints could be reliably differentiated, supervised machine learning (ML) models were applied to classify the AA-AuNPs spectra. **Figure 3** summarizes the classification performance of the RF model, which achieved a 100% accuracy on the test dataset, outperforming the SVM model that achieved 98.81% accuracy. The confusion matrix shown in panel **Figure 3a** demonstrates accurate classification of all amino acid-derived nanotags based on their spectral signatures. Furthermore, the two-dimensional t-SNE projection shown in **Figure S14a** reveals distinct clustering of the spectra, confirming that the biomineralized amino acid shells generate highly separable spectra populations. To further assess inter-class spectral relationships, a pairwise correlation heatmap was constructed across all seven AA-AuNP formulations **(Figure S14b).** Proline exhibited the lowest pairwise correlation with the remaining amino acids, consistent with its unique cyclic side-chain structure and the comparatively higher spectral variance observed in this nanotag class, while glutamic acid, serine, leucine, and valine showed stronger mutual correlations, reflecting their closer spectral profiles The mean spectra for each AA-AuNP, presented with standard deviation in **Figure 3b**. Among the seven amino acids, Pro-AuNPs exhibited comparatively larger spectral variance, which may contribute to the minor classification ambiguity observed in the confusion matrix in panel Figure 3a. To better understand the spectral features governing classification, RF feature importance analysis was performed to identify the top 20 discriminative wavenumbers (**Figure 3d**). Comparison of these wavenumbers with the averaged spectra (**Figure 3c**) demonstrated strong agreement between the most important classification features and the dominant Raman peaks, enabling an interpretable classification framework rather than a purely black-box prediction model. To complement the RF feature importance analysis, SHAP-based interpretation was applied to identify and determine the contribution of individual wavenumbers to the AA-AuNPs classification outcome. The top 10 SHAP-identified discriminative wavenumbers overlaid onto the average SERS spectra are shown in **Figure S15a**, with the corresponding normalized SHAP importance profiles presented in **Figure S15b**. This corroborated the RF feature importance analysis results, with the most discriminative wavenumbers concentrated in spectral regions corresponding to the dominant IR-780 Raman peaks. Notably, the three most significant wavenumbers for classification-530.0, 1204.0, and 1366.0 cm^-1^,corresponded to high-intensity spectral features with minimal variance across replicates.

**Figure 3.**
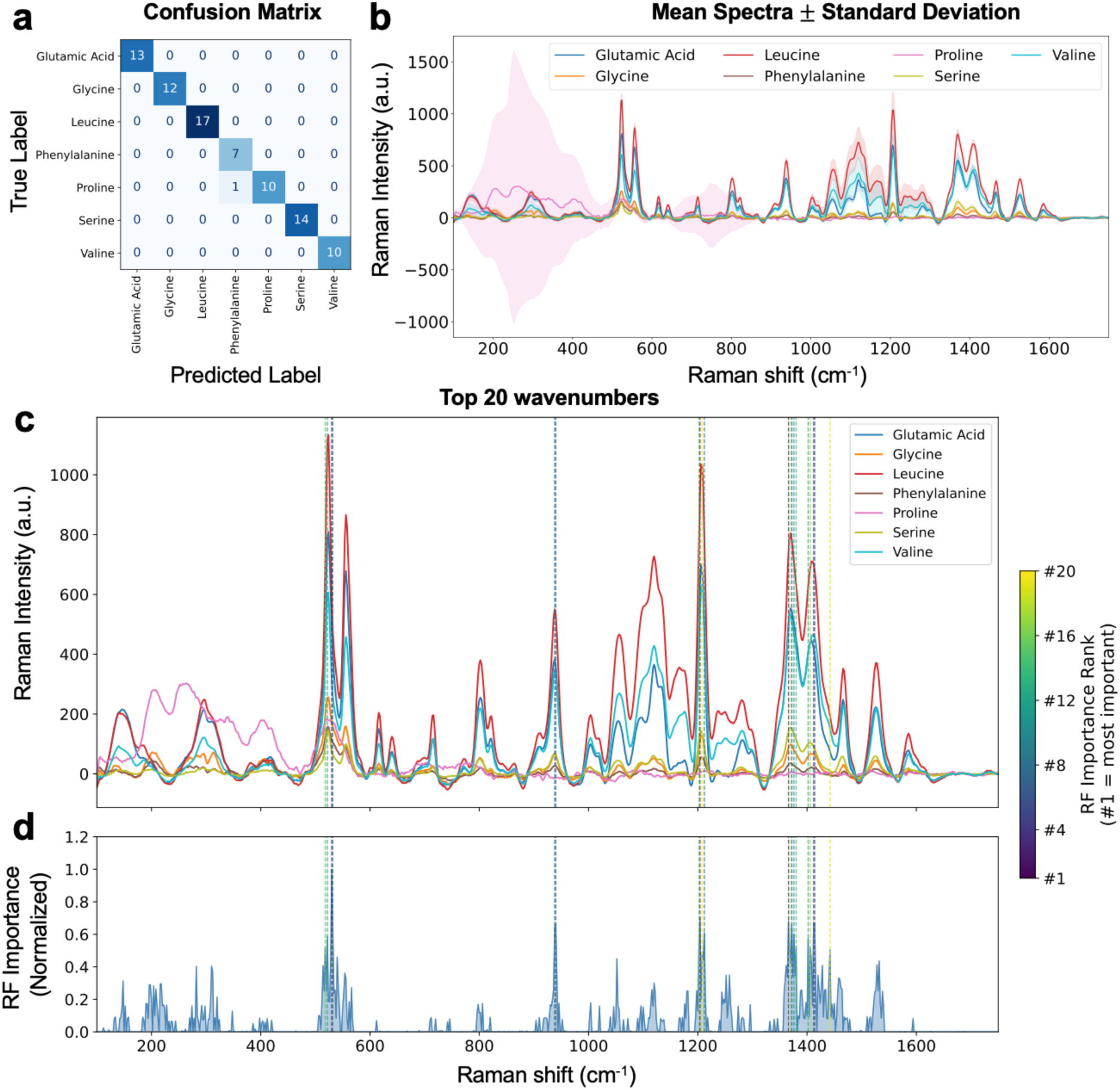
Machine learning-based classification of AA-AuNPs SERS spectra using an Random Forest model. (a) Confusion matrix showing classification performance on the test dataset. (b) Mean SERS spectra with standard deviation for each AA-AuNPs nanotags (c) Average SERS spectra overlaid with the top 20 discriminative wavenumbers identified by RF feature importance analysis, where dashed lines are color-coded according to feature rank. (d) Normalized RF feature importance profile across the spectral range, highlighting the relative contribution of the most informative Raman shifts used for classification.

To further evaluate model robustness, the trained RF classifier was applied to a completely independent single-blind test dataset. The model successfully classified 79 out of 80 AA-AuNP spectra, achieving 98.75% accuracy and demonstrating strong reproducibility across independent experiments **(Figure S16).** The single misclassification, where leucine was predicted as valine, is likely attributable to the close spectral similarity between these amino acids, consistent with their proximity in the t-SNE clustering analysis shown in **Figure S14a.**

### Ratiometric Differentiation and Mixed-Signature Resolution between Two AA-AuNPs

While the classification of discrete AA-AuNPs populations demonstrates that biomineralized nanotags generate chemically distinguishable SERS fingerprints, biological systems often contain heterogeneous or partially overlapping molecular environments rather than isolated spectral states.^10^ To evaluate whether the ML framework could resolve gradual compositional changes and mixed spectral signatures, we next investigated ratiometric mixtures of proline- and valine-derived AA-AuNPs across multiple concentration ratios.

Figure 4 summarizes the performance of the ML models for ratiometric classification of mixed Proline:Valine AA-AuNP spectra. Both ML classifiers (RF and SVM) achieved 100% accuracy on the test dataset, as demonstrated by the confusion matrix in panel **Figure 4a**, where all five compositional classes (P0V10, P10V0, P3V7, P5V5, P7V3) were correctly identified. The high classification accuracy suggests that the ratiometric mixture spectra retained sufficiently distinct and reproducible compositional signatures for reliable discrimination by both ML models. The mean SERS spectra (± standard deviation) for each Pro:Val AA-AuNP ratio are shown in **Figure 4b**. A gradual spectral evolution is observed across the compositional series, with the spectral profiles transitioning systematically between pure Val-AuNP and pure Pro-AuNP nanotag populations, demonstrating that mixed AA-AuNP systems generate composition-dependent SERS fingerprints. RF feature importance analysis revealed that the most discriminative spectral regions were concentrated primarily within the 1000–1600 cm^-1^ range, as shown in the normalized importance profile in **Figure 4e**.

**Figure 4.**
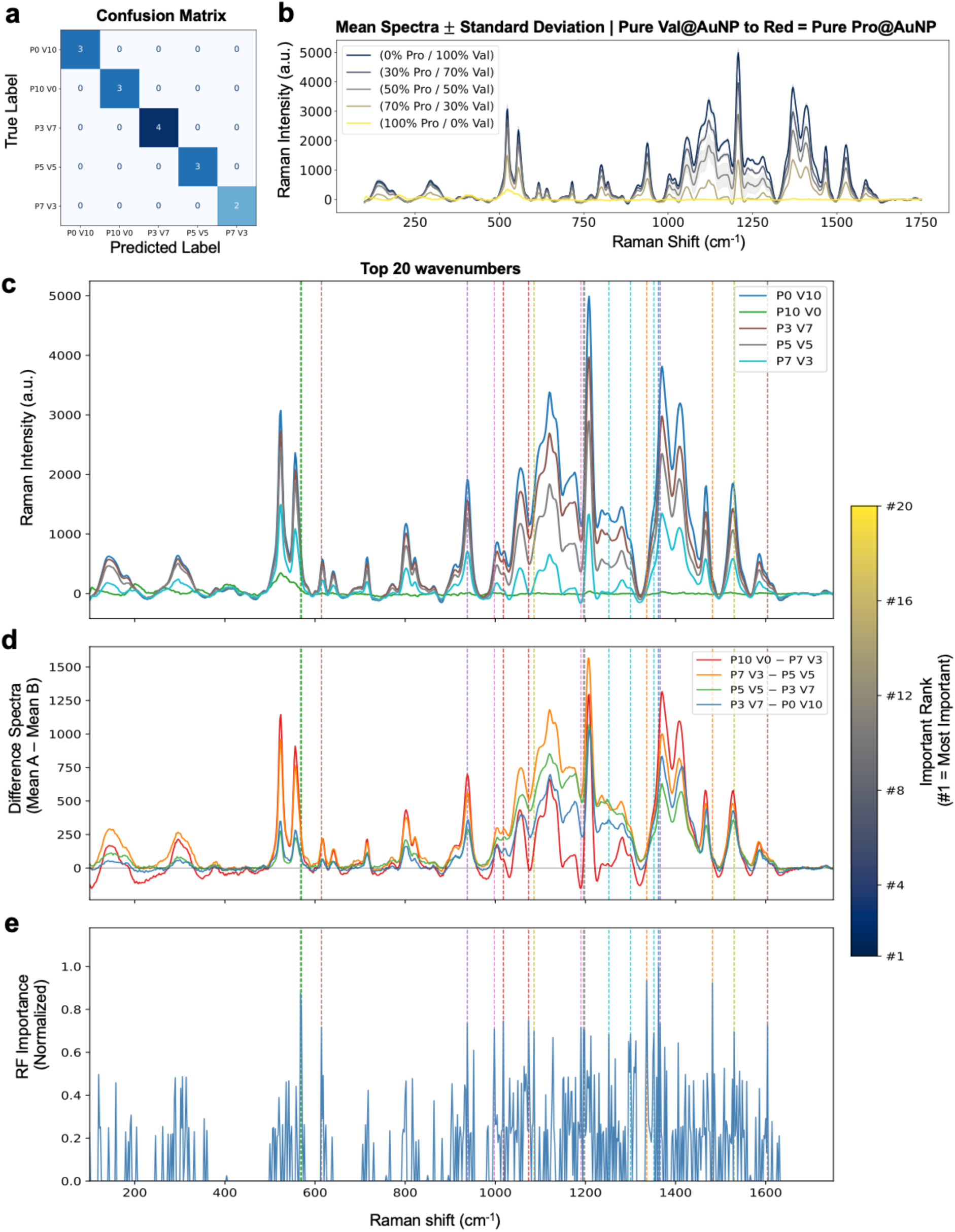
Ratiometric classification of Pro:Val AA-AuNP mixtures using a Random Forest (RF) model. (a) Confusion matrix showing classification performance on the test dataset. (b) Mean SERS spectra (± standard deviation) for each Pro:Val compositional ratio. (c) Averaged spectra overlaid with the top 20 discriminative Raman shifts identified by RF feature importance analysis, where dashed lines are color-coded according to feature rank. (d) Pairwise difference spectra (Mean A − Mean B) highlighting spectral regions contributing to class separation. (e) Normalized RF feature importance profile across the spectral range used for classification.

To interpret the spectral basis of classification, the top 20 most important Raman shifts identified by the RF model were overlaid onto the averaged spectra (**Figure 4c**), with dashed lines color-coded according to feature rank. Comparison with the pairwise difference spectra (Mean A − Mean B) shown in **Figure 4d** demonstrated that the RF model preferentially selected spectral regions exhibiting the greatest compositional variation between classes rather than simply the highest-intensity Raman peaks. In addition to RF, SHAP-based feature importance analysis for the Pro: Val ratiometric classification was performed, which corroborates the discriminative spectral regions identified by the RF model **(Figure S17)**. These findings indicate that discriminative information is encoded primarily in differential intensity patterns and relative spectral evolution across the mixture series, highlighting the sensitivity of the ML framework for resolving subtle compositional differences in biomineralized SERS nanotags. Additionally, representative SERS spectra of a Glu: Ser ratiometric series further demonstrated composition-dependent spectral evolution across varying mixture ratios, consistent with the findings for the Pro:Val system **(Figure S18).**

### Biomineralization Driven by Pediatric Tumor Cell-Derived Exosomes and ML-Assisted Differentiation for Diagnostic Classification

While the AA-AuNPs system provides a highly controlled environment to study how individual molecular side chains dictate SERS fingerprints, tumor-derived exosomes present a significantly more complex challenge. In the AA-AuNP model, the surface chemistry is defined by a single, known amino acid species; in contrast, the exosomal surface comprises a heterogeneous proteomic corona. To determine if our biomineralized approach could navigate this biological complexity, we transitioned from these ‘clean’ synthetic templates to the multifaceted protein landscapes of cancer-derived vesicles, utilizing their innate biochemical diversity as a driver for lineage-specific diagnosis.^49–51^

The transition to an exosome-based biomineralization strategy was motivated by the need for a “self-reporting” diagnostic platform that captures the complex tumor-associated biochemical states without exogenous labeling. To evaluate the translational potential of this approach, we systematically selected four clinically relevant pediatric tumor models representing two distinct developmental lineages: neuroblastoma (SK-N-BE(2) and COG-N-519) from neural crest-derived progenitors, and osteosarcoma (U2OS and OS-526), originating from mesenchymal osteoblast precursors (**Figure 5a**). Importantly, the neuroblastoma models were established from patient-derived pediatric tumors, enabling assessment of the SERS-ML platform in biologically heterogeneous and clinically relevant systems. This experimental design allowed us to determine whether biomineralized exosomal nano-bio interfaces retain sufficient lineage-specific molecular information to discriminate between tumor origins. Successful exosome isolation was confirmed by TEM imaging, which demonstrated intact lipid bilayers and spherical vesicular morphologies (**Figure 5b–e**), preserving the membrane-associated proteins and lipid architectures that govern the subsequent biomineralization process. ^52, 53^ BCA assay result confirms the presence of the protein within the isolated exosomes, as evidenced by the purple calorimetric response observed across all four cell line derived exosome samples **(Figure 5F, S19a).** This is consistent with the positive control of albumin standard of 2000 µg/mL and the negative control of PBS 1X, validating that the measured protein signal originates from exosomal protein content rather than background interference. SDS-PAGE analysis of exosomes isolated from those four cell lines reveals distinct protein banding patterns spanning ∼15–150 kDa molecular weight range, consistent with the canonical proteomic signature of exosomes, which characteristically co-enrich tetraspanins (CD9, CD63, CD81; ∼24–60 kDa) and heat shock proteins (Hsp70, ∼70 kDa) alongside MVB-associated proteins such as Alix and TSG101**(Figure 5g, S19b)**.^54–56^.

**Figure 5.**
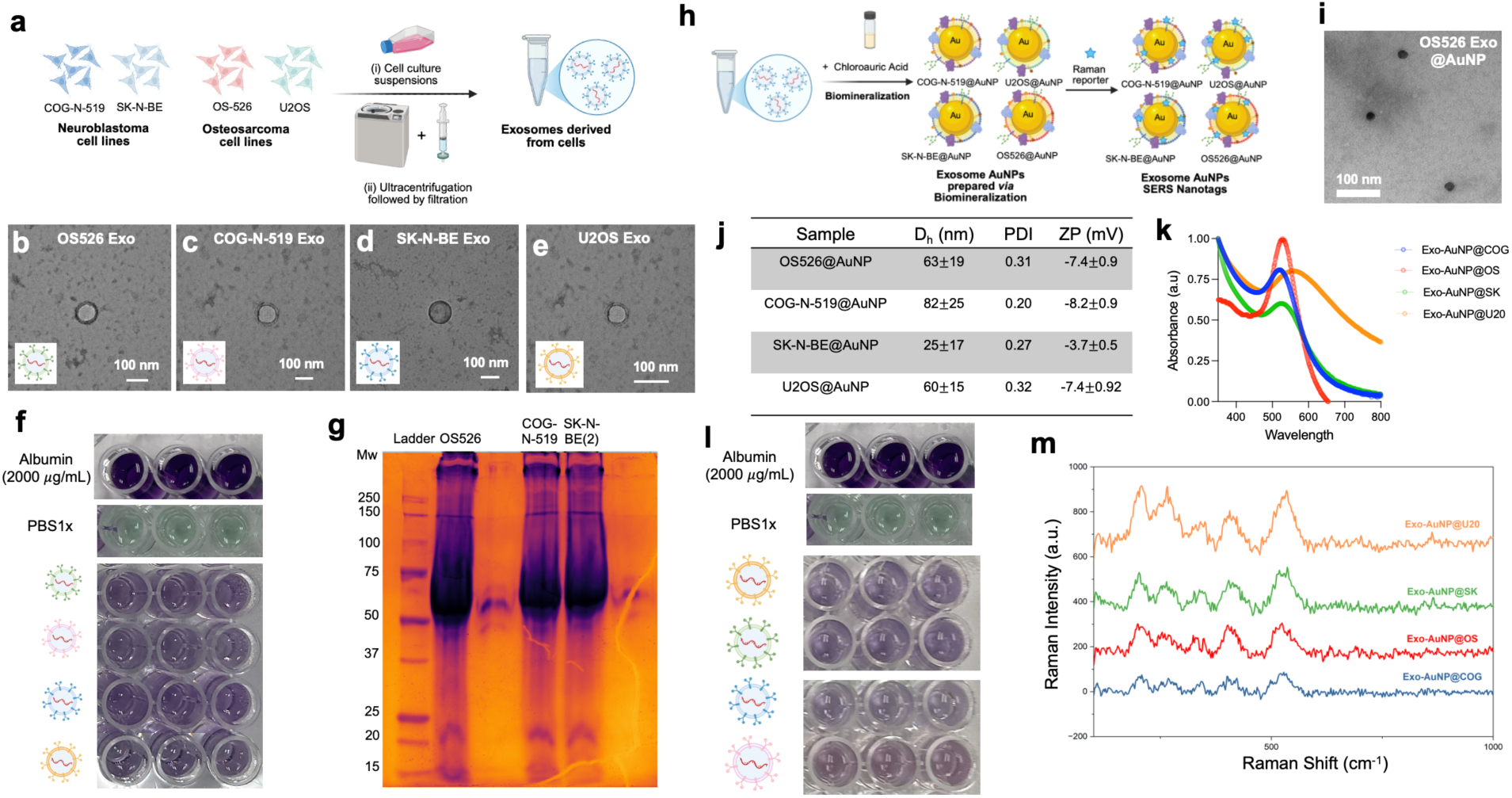
Isolation of exosomes from cell culture supernatants, synthesis and characterization of exosome biomineralized AuNPs (Exo-AuNPs). (a) Schematic of exosome isolation from COG-N-519, SK-N-BE (2), OS-526, U2OS cell culture supernatants using the ultra-centrifugation process. TEM image of exosome isolated from (b) OS-526 cell culture supernatant, (c) COG-N-519 cell culture supernatant, (d) SK-N-BE (2) cell culture supernatant, (e) U2OS cell culture supernatant. (f) BCA assay demonstrating the protein concentrations of the isolated exosomes, (g) SDS-PAGE gel studies of the isolated exosomes. (h) Schematic of biomineralization of distinct AuNPs using isolated exosomes to form Exo-AuNPs nanotags. (i) TEM image of spherical and highly uniform OS-526 Exo-AuNPs. (j) Physiochemical properties of the Exo-AuNPs including their hydrodynamic diameter, polydispersity index and Zeta potential, (k) UV-Vis absorbance. (l) BCA assay confirming presence of protein after biomineralization in Exo-AuNPs, and (m) distinct SERS spectra of the Exo-AuNPs.

The synthesis of Exo-AuNPs was achieved through a biomimetic reduction process in which the exosomal proteome serves as a multifunctional reactor. Upon the addition of isolated exosomes to the gold precursor under mild alkaline conditions, the biomineralization process is governed by a competitive interplay of specific amino acid residues within the protein corona. Specifically, the redox activity of tyrosine residues facilitates the reduction of Au^3+^ to Au^0^ via deprotonation of the phenolic hydroxyl group. Simultaneously, cysteine residues provide robust colloidal stability by forming covalent Au–S coordination bonds, effectively passivating the nascent gold surface. These biomineralized exosome-capped gold nanoparticles produced distinct calorimetric profiles **(Figure S20a)** and their TEM analysis revealed well-dispersed, spherical nanoparticles with a uniform, monodisperse size distribution (**Figure 5i).** Interestingly, the resulting Exo-AuNPs (25–83 nm) were consistently smaller than the source vesicles. This suggests a localized nucleation model, where particle growth occurs at discrete molecular “hot spots” defined by the density of tyrosine and cysteine residues, rather than a global reduction across the entire exosomal surface.^29^

The colloidal stability of Exo-AuNPs represents a marked departure from the primarily electrostatic stabilization observed in synthetic AA-AuNP systems. Here, stability is governed predominantly by steric hindrance arising from the dense biomolecular shell composed of membrane-associated proteins and lipids retained during biomineralization. TEM analysis confirmed the formation of spherical and highly uniform Exo-AuNPs (**Figure S20b**), while physicochemical characterization demonstrated distinct hydrodynamic diameters, polydispersity indices, and zeta potentials across the different exosome-derived formulations (**Figure 5j)**. UV–Vis spectroscopy further revealed distinct plasmonic absorption profiles for each Exo-AuNP population (**Figure 5k**), suggesting that lineage-dependent exosomal composition influences nanoparticle growth and optical behavior. Retention of exosome-associated proteins following biomineralization was confirmed by BCA analysis (**Figure 5l, S21**), supporting preservation of the native biomolecular interface after AuNP formation. The ultimate diagnostic readout was provided by the intrinsic SERS activity of the Exo-AuNPs (**Figure 5m, S22**).

Unlike the AA-AuNP systems, where reporter organization is governed primarily by a single amino acid-defined interface, the Exo-AuNPs present highly heterogeneous nano-bio architectures composed of membrane proteins, lipids, glycans, and associated biomolecular cargo. Consequently, the resulting SERS fingerprints likely emerge from collective variations in exosomal membrane composition, protein density, lipid organization, and biomolecular packing that modulate nanoparticle nucleation behavior, plasmonic hotspot formation, and reporter accessibility at the Au surface. Because exosomal membranes retain lineage-specific molecular signatures inherited from their parental tumor cells, the biomineralization process effectively encodes tumor-dependent biochemical information into the resulting plasmonic nanostructures. Variations in membrane-associated proteins and lipid microdomains may alter local electromagnetic coupling, hydrophobic reporter partitioning, steric accessibility, and reporter orientation within hotspot-rich regions, thereby generating distinct spectral morphologies across osteosarcoma-and neuroblastoma-derived Exo-AuNPs.

Importantly, the spectral behavior of Exo-AuNPs nanotags differed substantially from that observed in the amino acid-biomineralized systems. Whereas AA-AuNPs nanotags exhibited strong reporter-associated Raman features extending throughout the fingerprint region (1000–1600 cm^-1^), Exo-AuNP spectra were dominated primarily by low-wavenumber bands between 100–700 cm^-1^ with comparatively weak higher-wavenumber reporter features. This spectral redistribution likely arises from a combination of plasmonic, and interfacial effects associated with exosome-mediated biomineralization. In AA-AuNPs systems, the amino acid shell forms a relatively thin and chemically homogeneous interface that permits close association of IR-780 molecules with plasmonic hotspot regions on the Au surface, thereby enabling strong enhancement of characteristic reporter vibrations (**Figure 6a, c**).^47, 57^ In contrast, exosomal biomineralization possibly generates a chemically heterogeneous and structurally disordered nano-bio interfacial environments composed of membrane-associated proteins, lipids, glycans, and extracellular biomolecular cargo. This complex interfacial layer likely increases the average reporter-to-metal separation distance and alters reporter orientation, packing density, and accessibility to localized electromagnetic hotspots (**Figure 6b, c**). Additionally, the Exo-AuNPs were generally smaller than the AA-AuNPs, which may further reduce electromagnetic field enhancement because SERS intensity is strongly dependent on nanoparticle size, plasmonic coupling efficiency, and hotspot formation.^57, 58^ Because SERS enhancement decays exponentially with increasing distance from the plasmonic surface, these combined effects may selectively suppress higher-wavenumber IR-780 vibrational modes, causing the spectra to become increasingly dominated by low-wavenumber interfacial vibrations associated with Au-biomolecule coordination environments and sulfur-containing biomolecular modes (**Figure 6b, d**).^59, 60^ Consequently, the Exo-AuNP spectra appear to probe the biomineralized nano-bio interface itself rather than predominantly reflecting the intrinsic Raman fingerprint of the reporter molecule. This mechanistic distinction further supports the hypothesis that biomolecular identity is encoded into the SERS response through modulation of reporter organization and plasmonic coupling at the nanoparticle interface (**Figure 6d**). Collectively, these findings suggest that biomineralization does not merely preserve exosomal biochemical identity but actively transforms complex membrane-associated molecular information into distinct lineage-dependent SERS fingerprints suitable for diagnostic classification.

**Figure 6.**
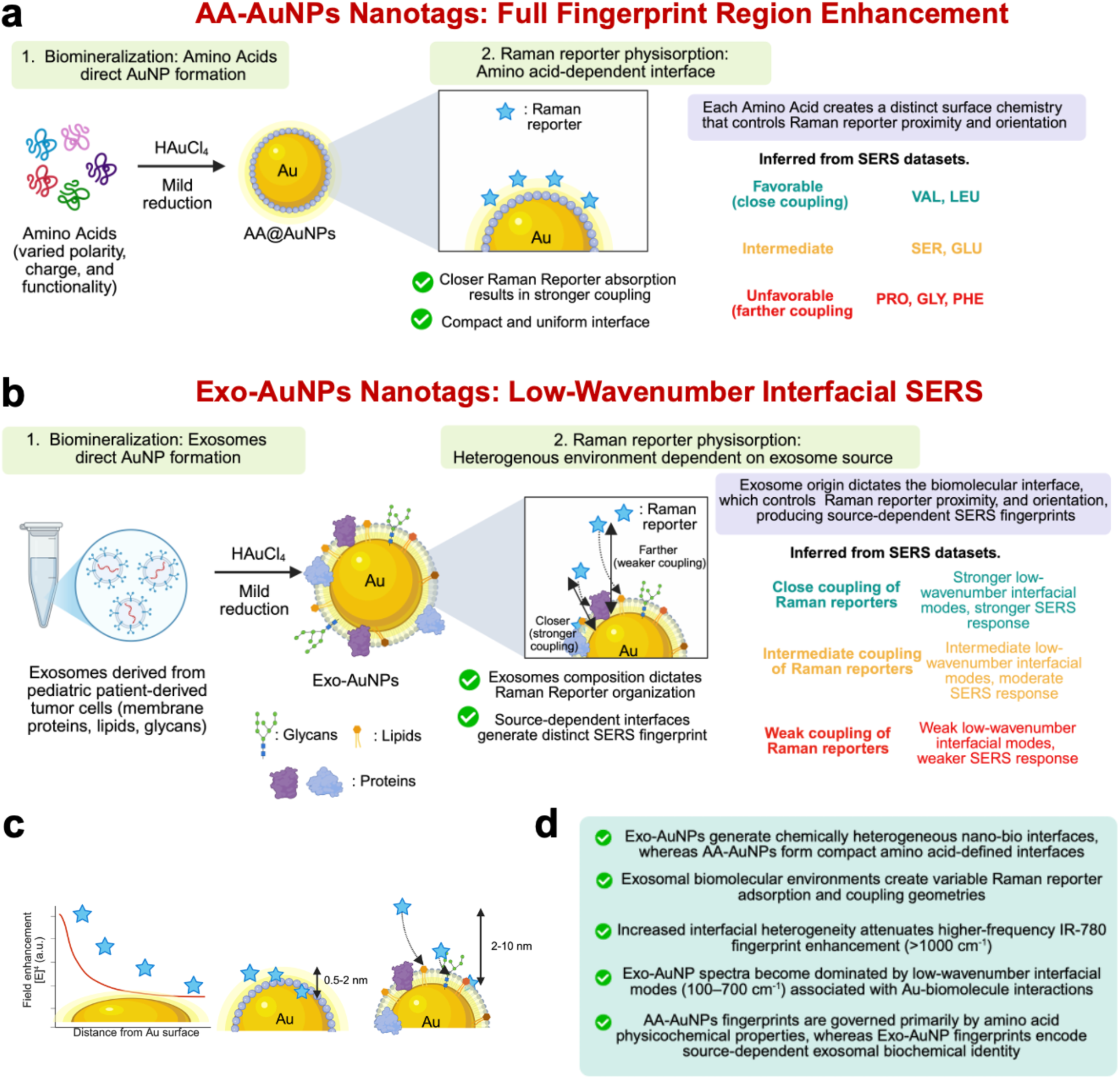
Proposed mechanistic model describing biomineralization-dependent modulation of SERS fingerprints in AA-AuNPs and Exo-AuNPs. (a) Amino acid-mediated biomineralization generates chemically defined nano-bio interfaces that regulate IR-780 reporter proximity and plasmonic coupling, producing variable enhancement across the full Raman fingerprint region (400-1600 cm^-1^). Favorable amino acid environments promote closer reporter–metal coupling and stronger reporter-associated vibrational enhancement. (b) Exosome-mediated biomineralization generates chemically heterogeneous nano-bio interfacial environments composed of membrane-associated biomolecules that modulate reporter organization, adsorption geometry, and hotspot accessibility, resulting in source-dependent SERS fingerprints dominated by low-wavenumber interfacial modes (100–700 cm^-1^). (c) Conceptual illustration of distance-dependent electromagnetic enhancement in SERS. Compact AA-AuNPs interfaces permit closer reporter adsorption and stronger near-field coupling, whereas heterogeneous Exo-AuNPs interfaces generate variable reporter–metal separation distances that attenuate higher-frequency IR-780 fingerprint enhancement. (d) Summary of the proposed mechanistic differences between AA-AuNPs and Exo-AuNPs and their resulting SERS spectral behavior.

### ML-Assisted Classification of Exosome-Biomineralized AuNPs

To determine whether exosome-derived biomineralized AuNPs encode tumor-specific biochemical information, we next applied supervised ML models to classify SERS spectra collected from Exo-AuNPs nanotags. Unlike the AA-AuNPs nanotags, where the surface chemistry is defined by a single biomolecular template, Exo-AuNPs present a more complex and heterogeneous protein corona. Therefore, successful classification in this setting would demonstrate that the SERS-ML platform can resolve biologically relevant spectral differences arising from tumor cell-derived extracellular vesicles.

Classification accuracy revealed clearer differences between the two models. RF achieved 83.33% accuracy, while SVM improved performance to 93.94% **(Figure 7a-e)**. The RF confusion matrix showed that misclassifications were primarily concentrated among the spectrally overlapping classes, including SK-N-BE(2), COG-N-519, OS-526, and U2OS. In contrast, SVM correctly classified all samples and reduced errors across the remaining classes, suggesting that its kernel- based decision boundary better captured nonlinear spectral differences within the exosome-derived dataset.

**Figure 7.**
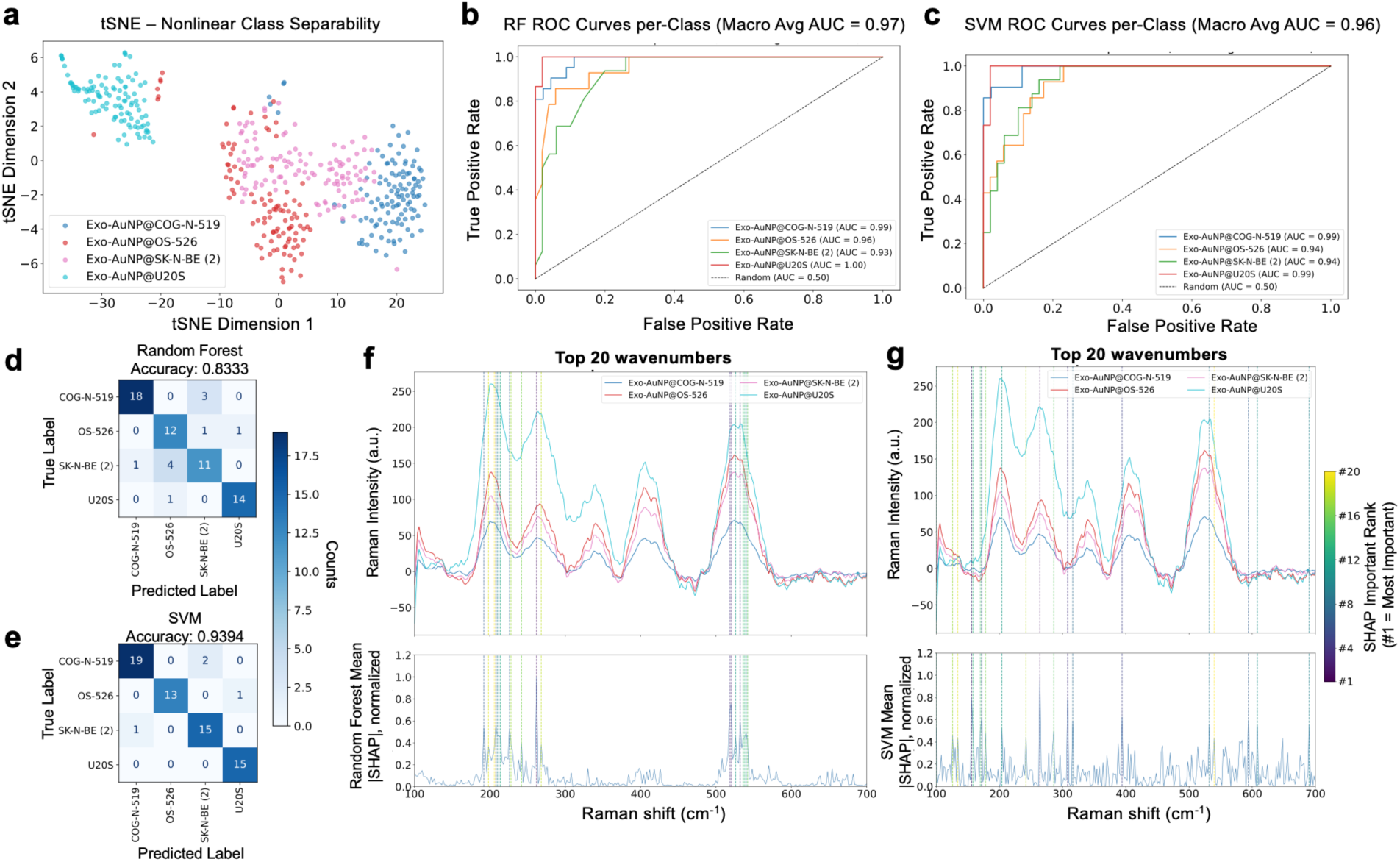
ML-assisted classification of SERS spectra collected from exosome-biomineralized AuNP nanotags (Exo-AuNPs). (a) t-SNE projection showing spectral clustering of Exo-AuNP nanotags derived from distinct pediatric tumor cell lines. (b, c) Receiver operating characteristic (ROC) curves for the RF and SVM models, respectively. (d, e) Confusion matrices showing classification performance of the RF and SVM models on the test dataset. (f, g) SHAP-based feature importance analysis for RF and SVM models, respectively, highlighting the most discriminative low-wavenumber spectral regions contributing to tumor classification.

To interpret the spectral basis of classification, SHAP-based feature importance analysis was performed for both models. Unlike conventional EV-SERS platforms, where intact extracellular vesicles are adsorbed onto pre-fabricated plasmonic substrates and generate broad biochemical signatures spanning proteins, lipids, nucleic acids, and glycocalyx-associated structures across the fingerprint region,^61,62^ our biomineralized Exo-AuNPs platform fundamentally probes the exosome-directed nanoparticle interface itself. Consequently, the most discriminative spectral information was concentrated within the low-wavenumber region (100–700 cm^-1^), as shown in **Figure 7f,g**. These low-wavenumber features are more sensitive to metal–biomolecule interfacial vibrations, including Au–S coordination environments and sulfur-containing protein modes associated with cysteine-rich exosomal surface proteins involved in biomineralization. In particular, bands within the 500–550 cm^-1^ and 640–650 cm^-1^ regions are consistent with C–S stretching and sulfur-associated protein vibrations, ^63, 64^ while features below 300 cm^-1^ may reflect Au–biomolecule coordination environments formed during exosome-mediated nanoparticle growth.^62, 65^

In contrast, higher-wavenumber lipid and nucleic acid vibrations commonly observed in substrate-based EV-SERS systems may be less pronounced in the present platform because plasmonic enhancement is localized primarily at the biomineralized protein corona rather than uniformly across the entire vesicular membrane architecture. RF feature importance remained concentrated within a relatively narrow subset of dominant interfacial features, whereas SVM distributed importance more broadly across the low-wavenumber region, suggesting that SVM leveraged subtle, distributed spectral variations that RF partially overlooked. This broader feature utilization likely contributed to the superior SVM classification accuracy in this biologically complex and partially overlapping dataset. Together, these findings demonstrate that exosome-driven biomineralization produces tumor-specific interfacial SERS fingerprints that can be resolved through ML-assisted analysis, supporting the potential of Exo-AuNP nanotags as a label-free diagnostic platform.

## CONCLUSIONS

In this study, we established a biomineralization-driven SERS platform that converts biomolecular identity into distinguishable plasmonic fingerprints for ML-assisted classification. By leveraging the intrinsic physicochemical properties of biomolecules as both reducing and stabilizing agents, we demonstrated that biomineralized AuNPs retain complex nano-bio interfacial chemistries capable of modulating Raman reporter organization, plasmonic hotspot accessibility, and electromagnetic coupling, thereby generating highly distinct SERS signatures. Using α-amino acids as a controlled proof-of-concept system, we showed that subtle differences in side-chain polarity, steric configuration, hydrophobicity, and electrostatic interactions produce distinct spectral fingerprints that can be classified with near-perfect accuracy using RF and SVM models. Furthermore, ratiometric analysis of mixed Pro-AuNP and Val-AuNP systems demonstrated that the platform is sensitive not only to discrete biomolecular states, but also to gradual compositional variation and mixed spectral populations.

Extending this approach to biologically complex systems, we synthesized Exo-AuNP nanotags using patient-derived pediatric tumor cell-derived exosomes from osteosarcoma and neuroblastoma models. Unlike conventional EV-SERS platforms that probe intact vesicles adsorbed onto external plasmonic substrates, our biomineralized approach directly interrogates exosome-mediated nano-bio interfaces composed of membrane proteins, lipids, glycans, and associated biomolecular cargo. Consequently, the resulting SERS fingerprints likely emerge from collective variations in exosomal membrane composition, protein density, lipid organization, and reporter accessibility within plasmonic hotspot-rich regions. Despite substantial biological heterogeneity and partial spectral overlap between tumor classes, SVM classification achieved 93.94% accuracy, while SHAP and t-SNE analyses enabled interpretable visualization of spectral clustering and discriminative Raman shifts.

Collectively, these findings demonstrate that biomineralization functions not only as a green synthetic strategy for plasmonic nanomaterials, but also as a mechanism for transducing molecular and cellular biochemical organization into high-dimensional optical fingerprints. By integrating biomineralized SERS nanotags with interpretable machine learning analysis, this work establishes a scalable and label-free framework for biomolecular sensing and tumor classification. More broadly, the ability to resolve biologically encoded interfacial spectral signatures highlights the potential of biomineralized plasmonic systems for future liquid biopsy diagnostics, precision oncology, and ML-enabled spectroscopic sensing applications.

## AUTHOR INFORMATION

## Author Contributions

I. S. conceived the idea and designed all the experiments. M.H.R. was involved in data collection and curation, formal analysis, investigation, methodology, validation, and visualization of all the experiments under the supervision of I.S. M.M. performed BCA assay and SDS PAGE under the supervision of I.S. T.N. performed mammalian cell culture experiments under the supervision of B. K. Exosomes collected *via* ultracentrifugation approach was collected by M.H.R. under the supervision of D.L. M.L. analysis were conducted by M.U.M. under the supervision of Z.A. The manuscript was written by I.S. with contribution from all the authors.

## ACKNOWLEDGMENT

I.S. acknowledges the Edward E. Whitacre Jr. College of Engineering and Texas Tech University for research support.

