## Supporting Information for "Biomineralized Surface-Enhanced Raman Scattering Nanotags Enable Machine Learning-Based Differentiation of Biomolecular Signatures"

### Biomineralized Surface-Enhanced Raman Scattering Nanotags Encode Biomolecular Identity into Machine Learning-Resolvable Plasmonic Fingerprints

*Md Hasnat Rashid,[1] Moin Uddin Maruf,[1] Troy Nations,[2] Mahenour Megahed,[1] Danielle E. Levitt,[3] Balakrishna Koneru,[2] Zeeshan Ahmad,[1] Indrajit Srivastava [1,4,]**

[1] Department of Mechanical Engineering, Edward E. Whitacre Jr. College of Engineering, Texas Tech University, Lubbock, Texas 79409, United States

[2] Pediatric Cancer Research Center, Department of Pediatrics, School of Medicine, Texas Tech University Health Sciences Center School of Medicine, Lubbock, Texas 79430, United States.

[3] Department of Kinesiology and Sport Management, Texas Tech University, Lubbock, Texas 79409, USA

[4] Texas Center for Comparative Cancer Research (TC3R), Amarillo, Texas, 79106, United States

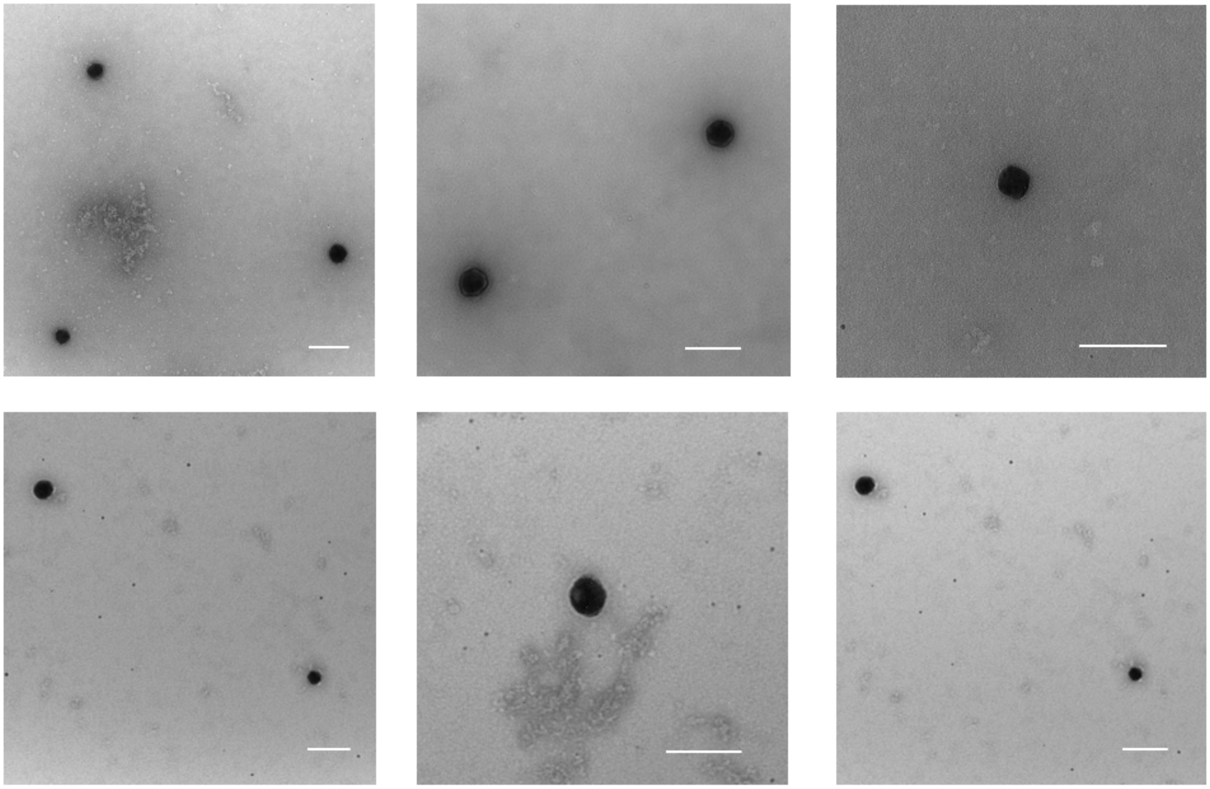

**Figure S1.** Representative negative-stained TEM images of Glu@AuNPs samples showing particle morphology (scale bar: 100nm)

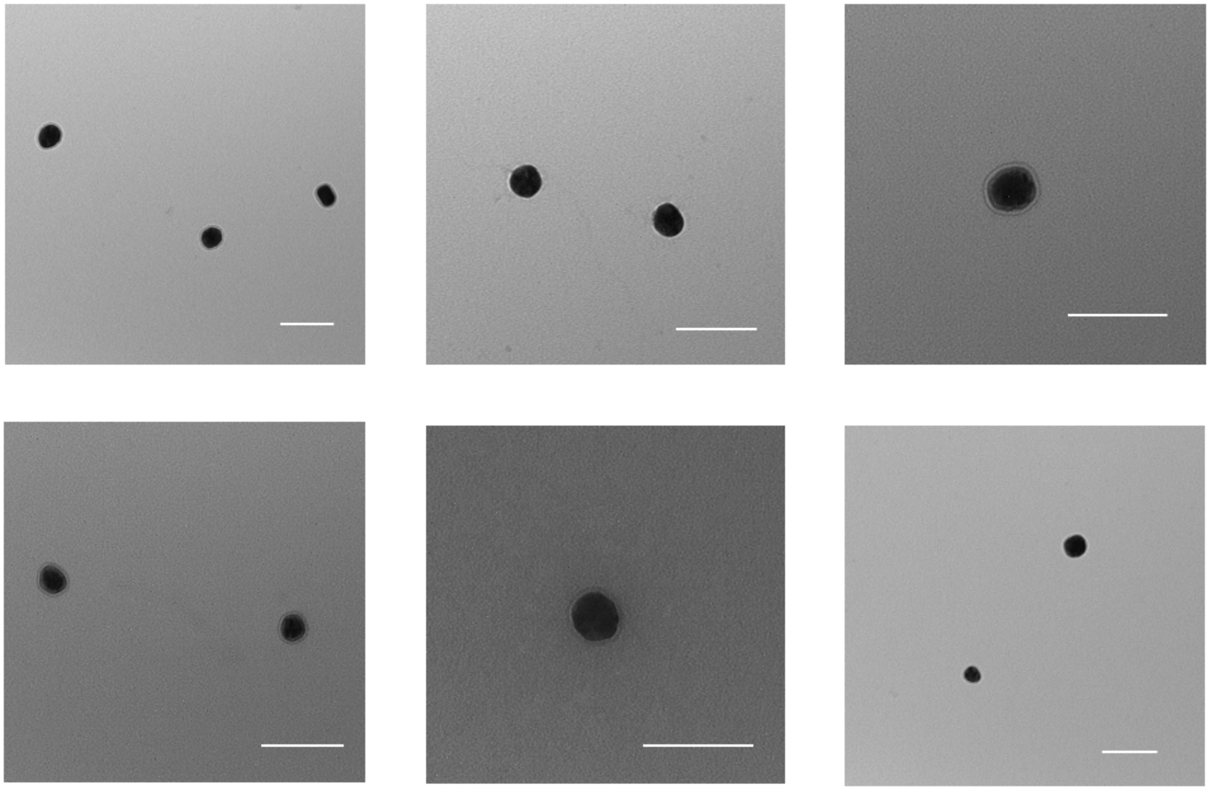

**Figure S2.** Representative negative-stained TEM images of Gly@AuNPs samples showing particle morphology (scale bar: 100nm)

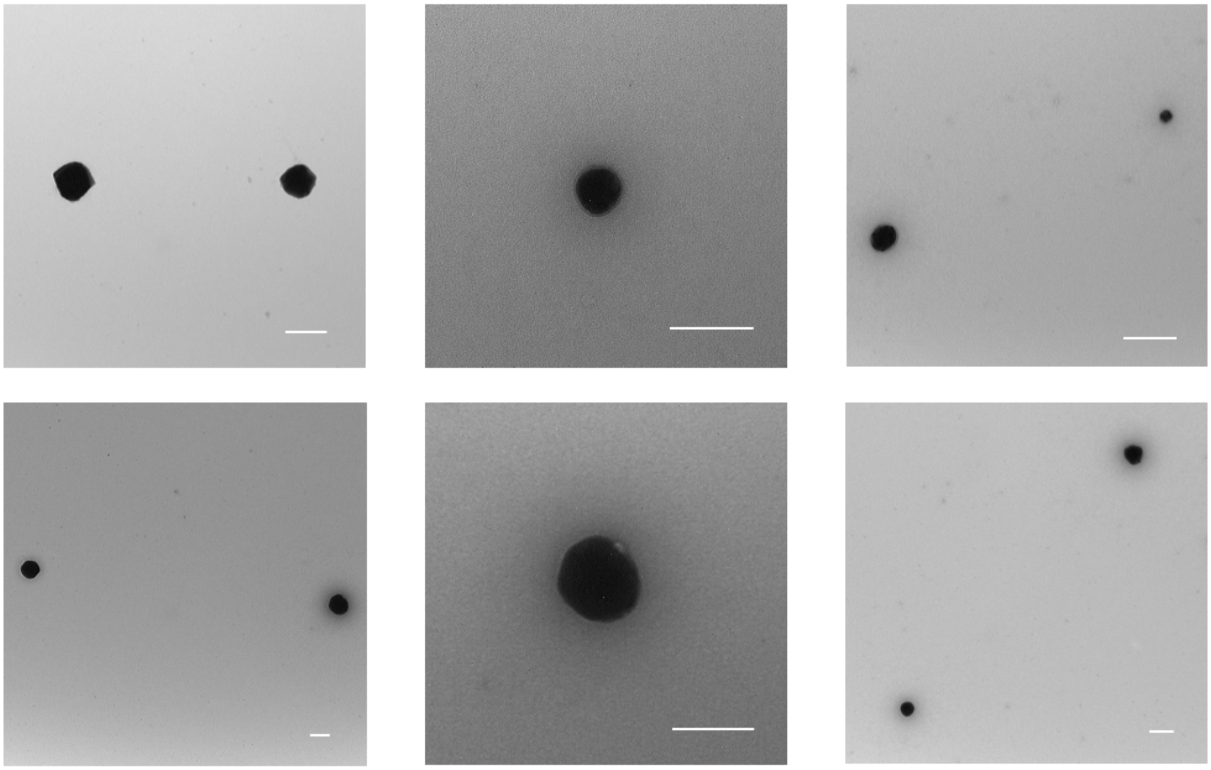

**Figure S3.** Representative negative-stained TEM images of Leu@AuNPs samples showing particle morphology (scale bar: 100nm)

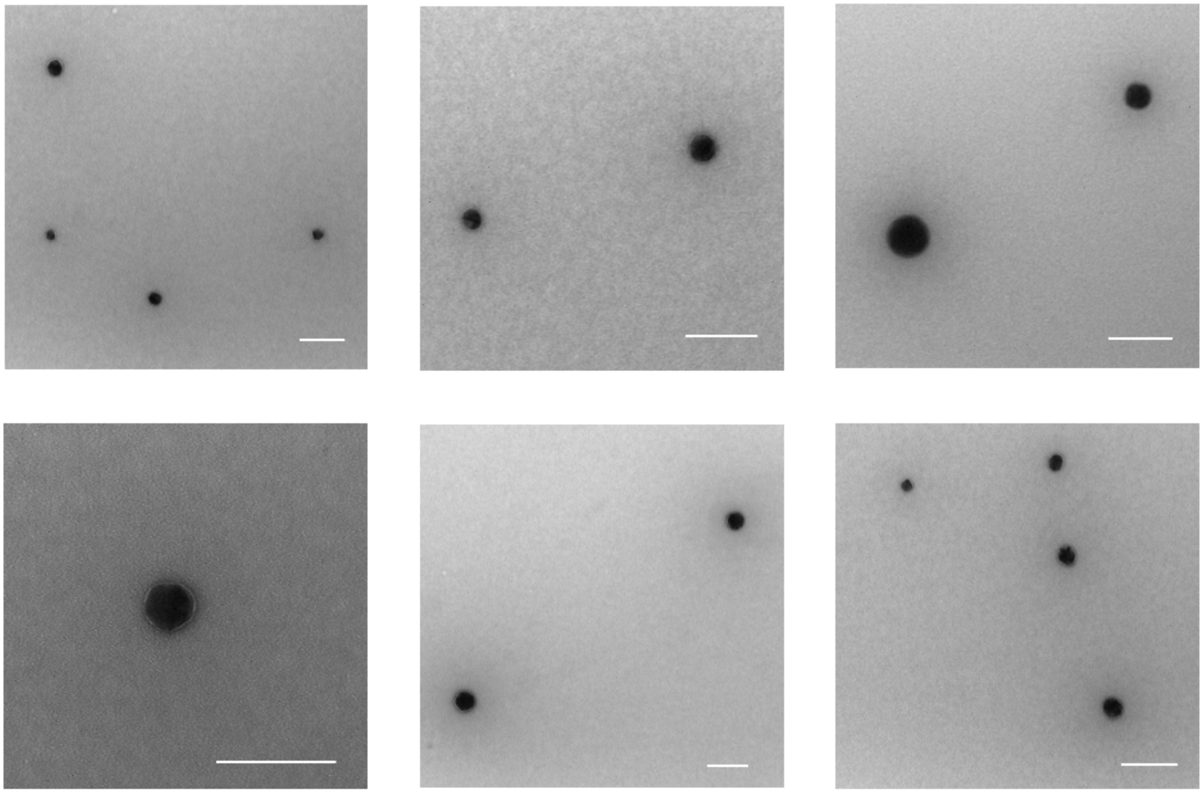

**Figure S4.** Representative negative-stained TEM images of Phe@AuNPs samples showing particle morphology (scale bar: 100nm)

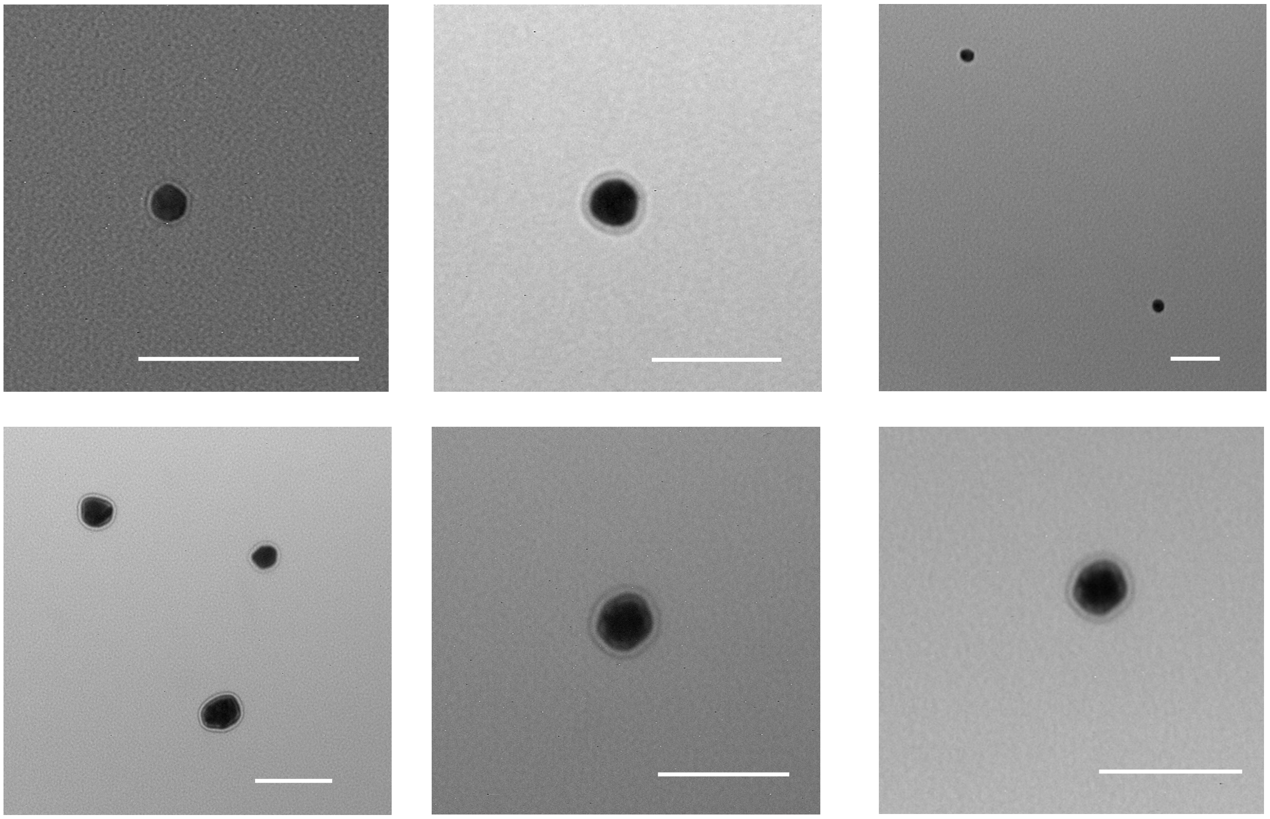

**Figure S5.** Representative negative-stained TEM images of Pro@AuNPs samples showing particle morphology (scale bar: 100nm)

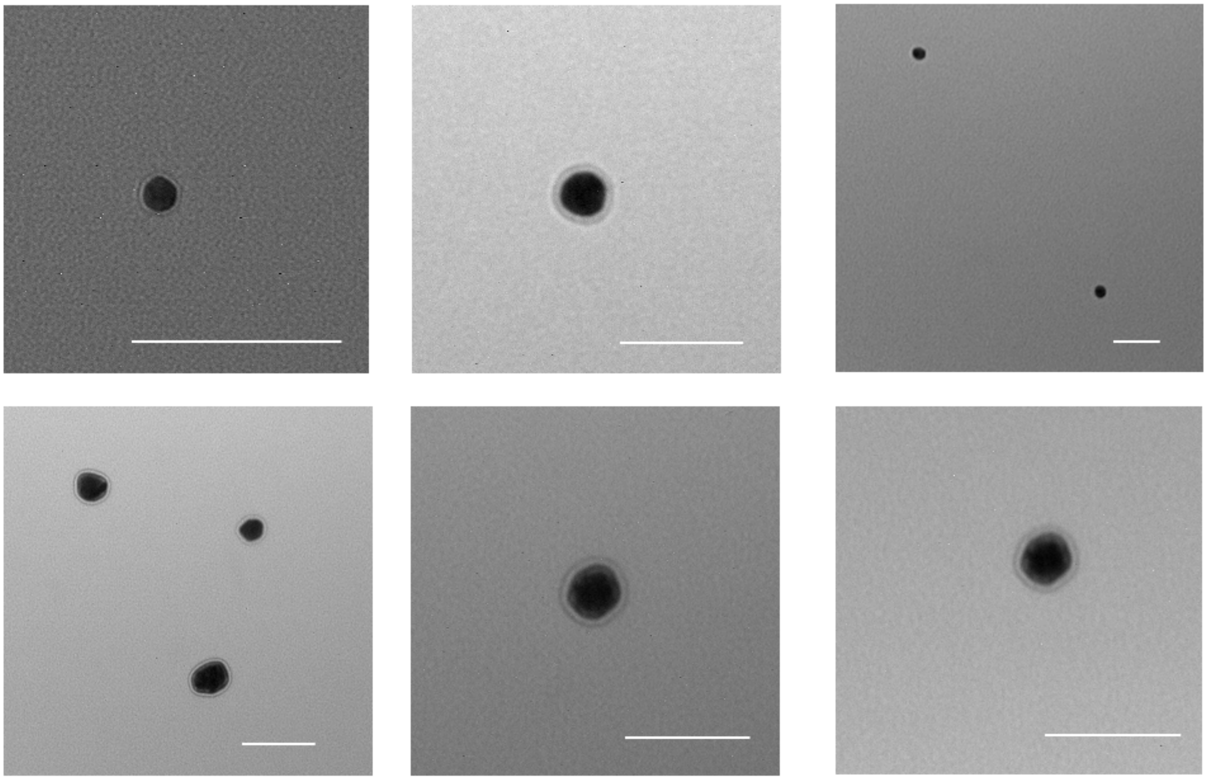

**Figure S6.** Representative negative-stained TEM images of Ser@AuNPs samples showing particle morphology (scale bar: 100nm)

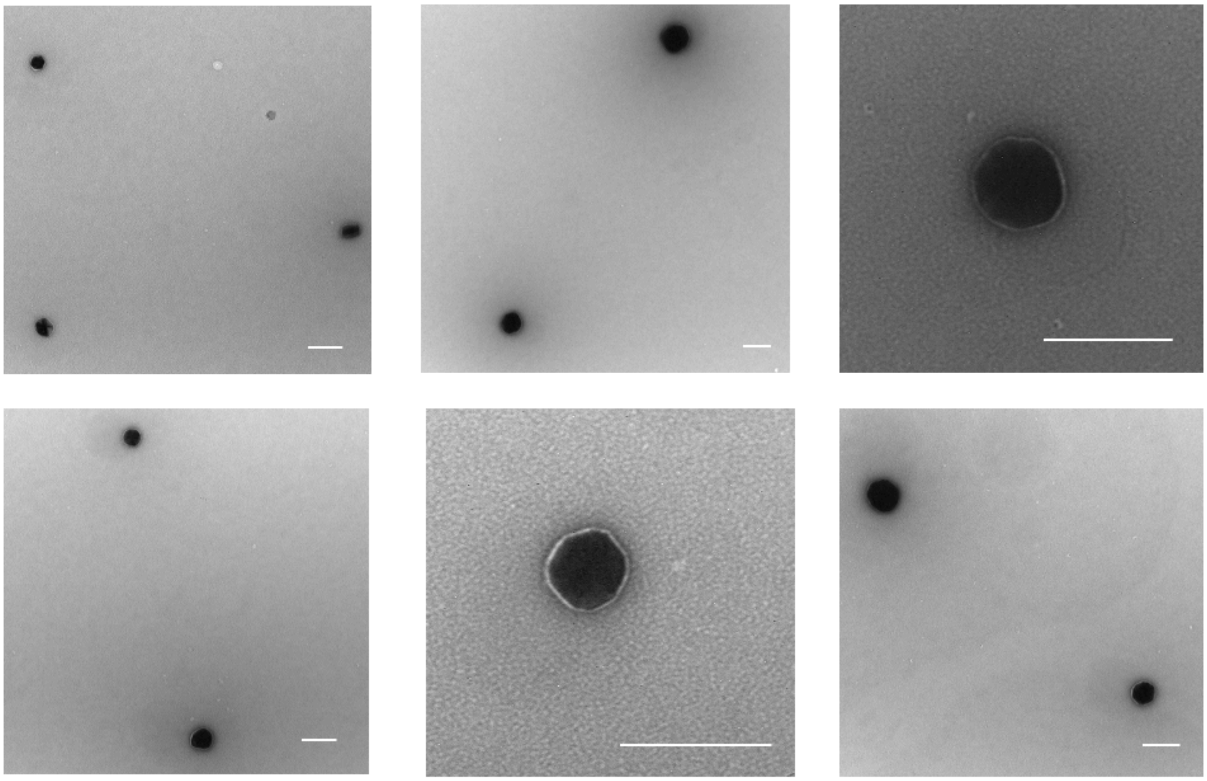

**Figure S7.** Representative negative-stained TEM images of Val@AuNPs samples showing particle morphology (scale bar: 100nm)

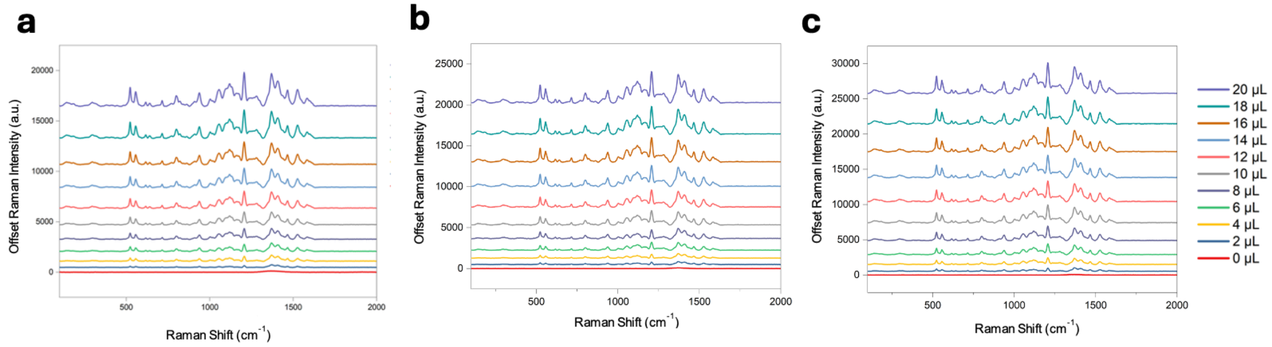


**Figure S8.** Stability of SERS spectra of Glu@AuNPs over three consecutive days. (a) Day 1, (b) Day 2, (c) Day 3.

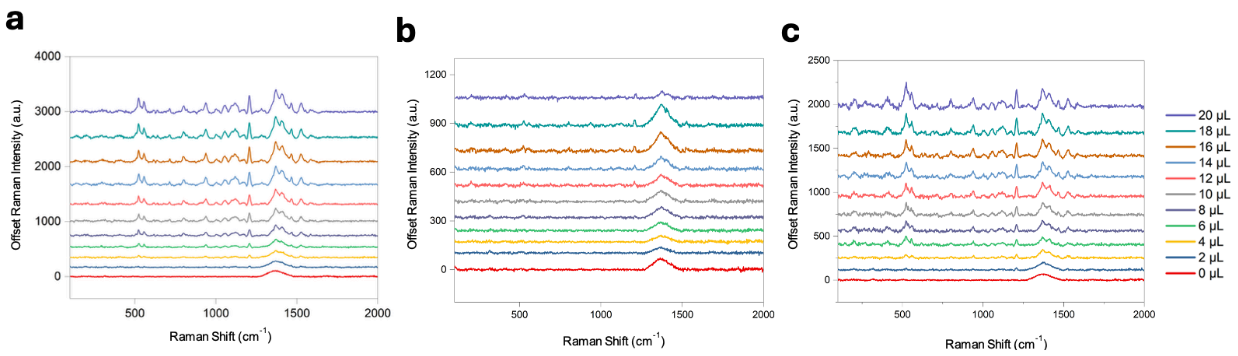


**Figure S9.** Stability of SERS spectra of Gly@AuNPs over three consecutive days. (a) Day 1, (b) Day 2, (c) Day 3.

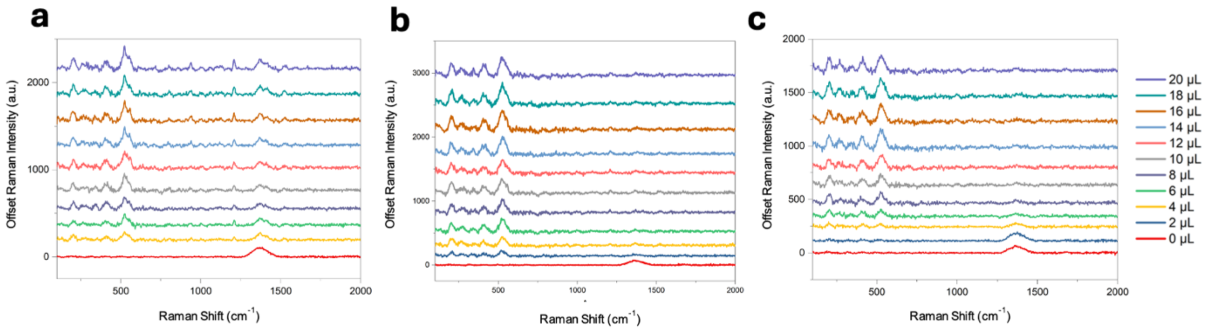


**Figure S10.** Stability of SERS spectra of Phe@AuNPs over three consecutive days, (a) Day 1, (b) Day 2, (c) Day 3.

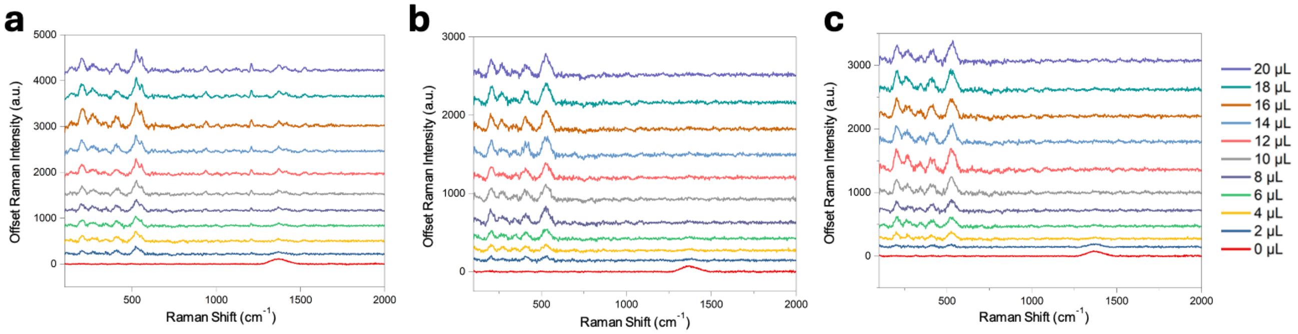


**Figure S11.** Stability of SERS spectra of Pro@AuNPs over three consecutive days, (a) Day 1, (b) Day 2, (c) Day 3.

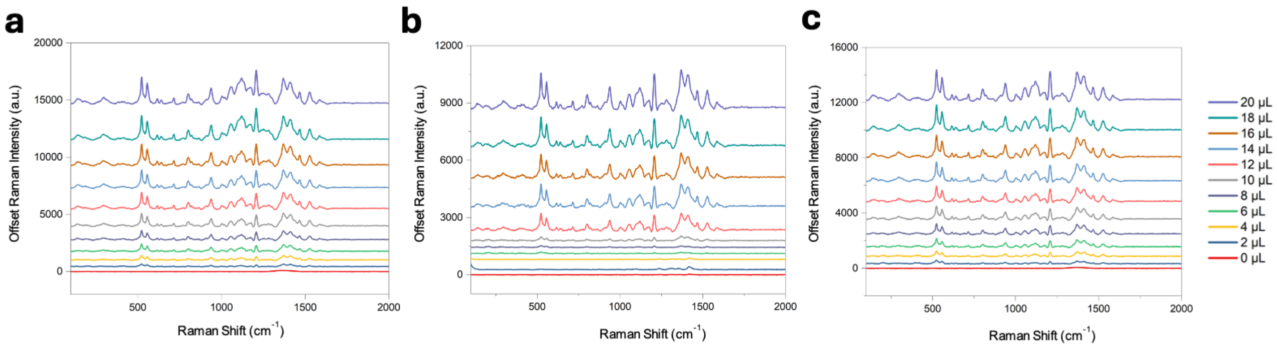


**Figure S12.** Stability of SERS spectra of Ser@AuNPs over three consecutive days, (a) Day 1, (b) Day 2, (c) Day 3.

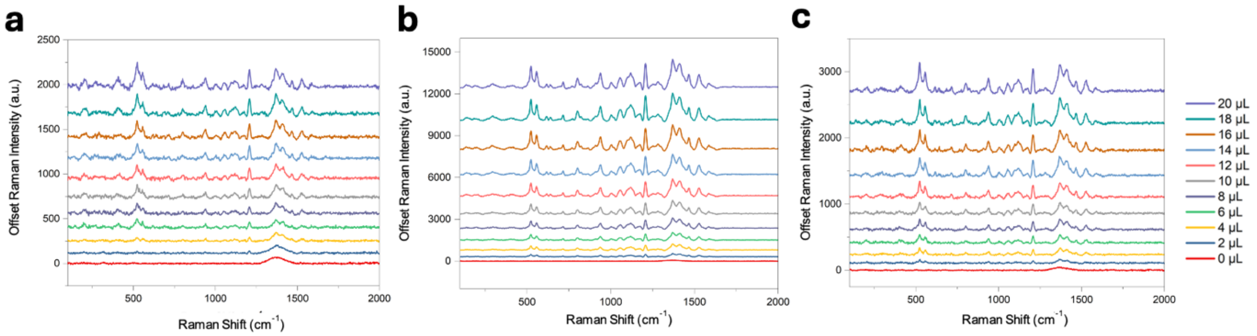


**Figure S13.** Stability of SERS spectra of Val@AuNPs over three consecutive days, (a) Day 1, (b) Day 2, (c) Day 3.

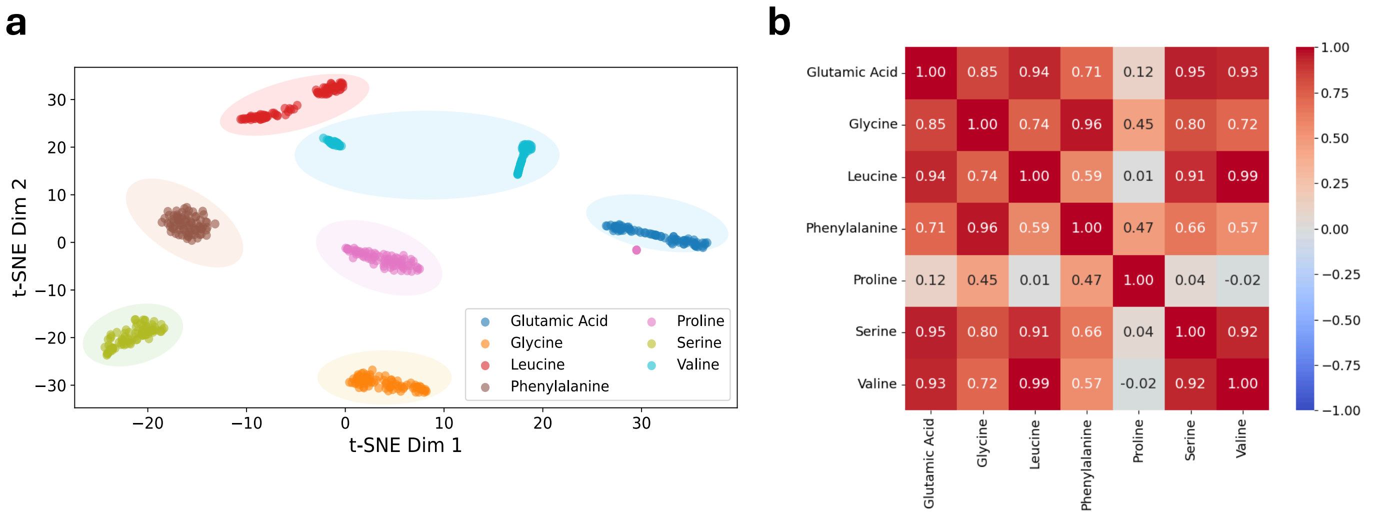


**Figure S14. Machine Learning Analysis of AA-AuNPs.** (a) t-SNE visualization showing clustering of different AA-AuNPs spectral profiles. (b) Pairwise spectral correlation heatmap between different AA-AuNPs.

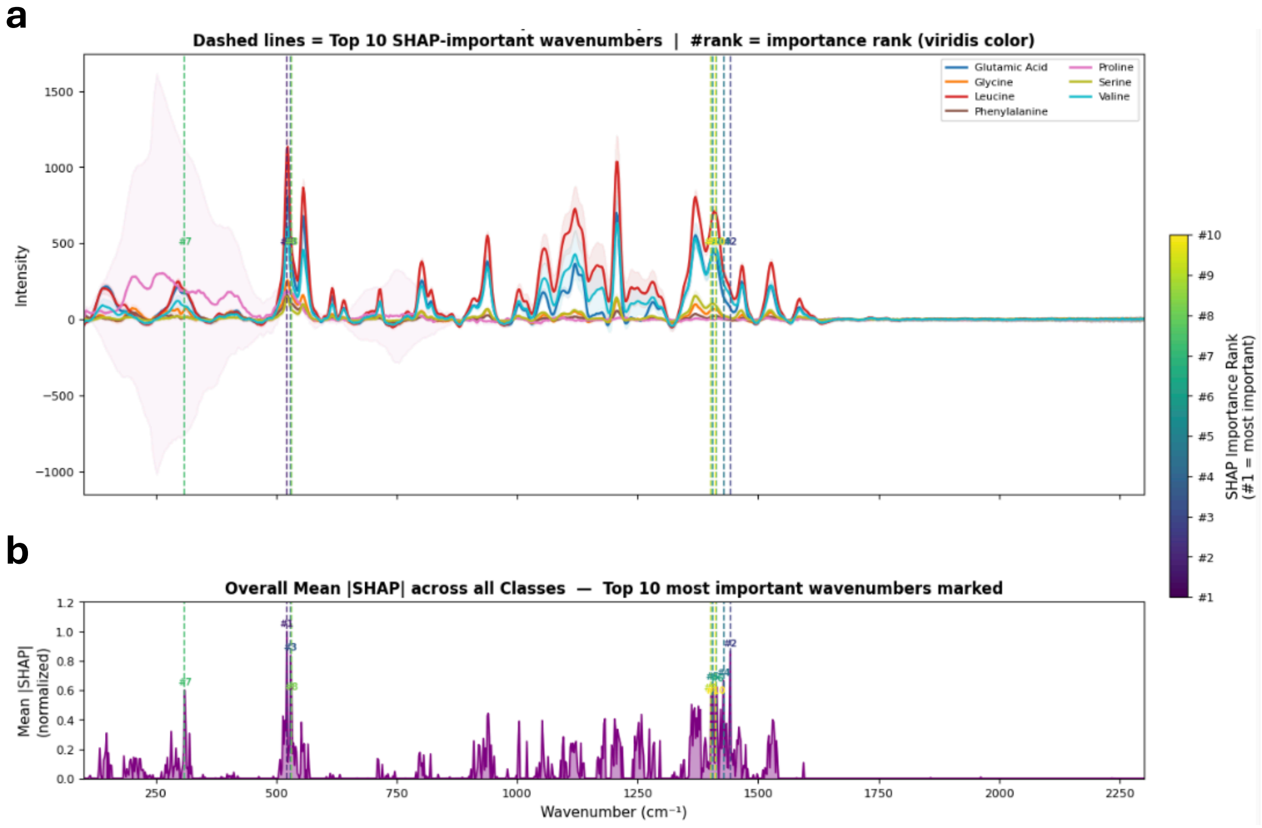


**Figure S15. SHAP feature importance analysis of AA-AuNPs.** (a) Average SERS spectra with the top 10 SHAP-identified discriminative wavenumbers, indicated by dashed lines color-coded according to importance rank. (b) Normalized SHAP feature importance profile highlighting key Raman shifts contributing to classification.

**a**

```
Prediction Summary (Counts of True vs Predicted):
Predicted  Glutamic Acid  Leucine  Phenylalanine  Serine  Valine
Label
AA1                    0       19              0       0       1
AA2                    0        0              0      20       0
AA3                    0        0             20       0       0
AA4                   20        0              0       0       0
```

**b**

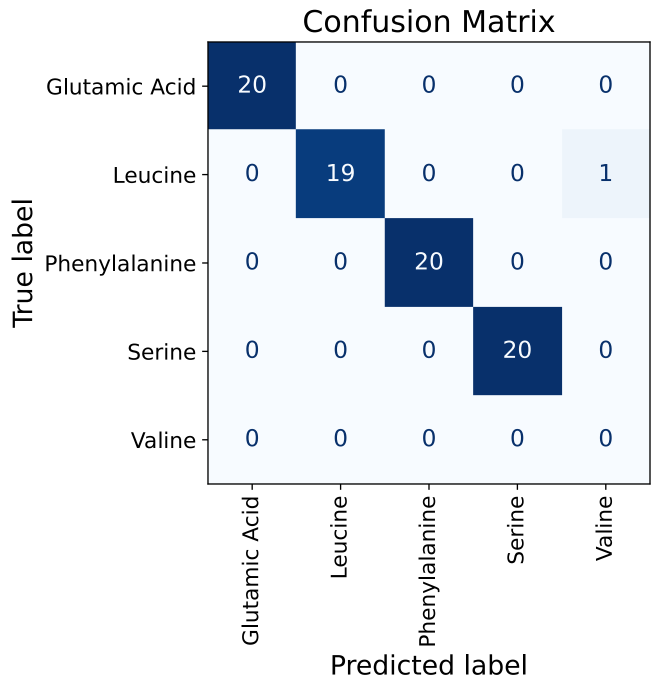


**Figure S16. Single-blind analysis for the identification of AA-AuNPs from mixed samples.** (a) Prediction summary of the Random Forest (RF) model showing true versus predicted classifications. (b) Confusion matrix illustrating the classification performance of the RF model in the single-blind study.

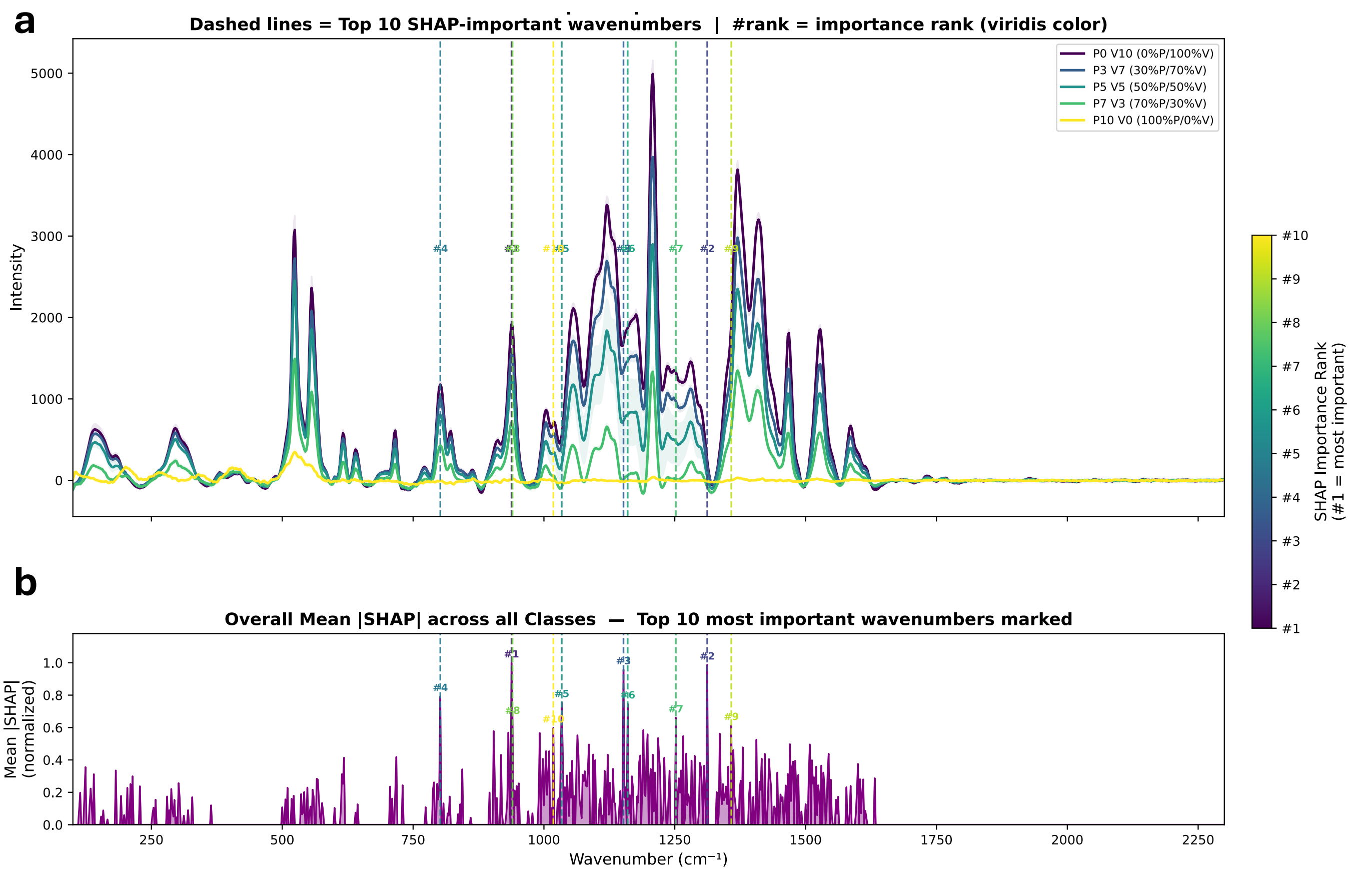


**Figure S17. SHAP feature importance analysis of Pro:Val AA-AuNPs ratiometric mixtures.** (a) Average SERS spectra showing the top 10 SHAP-identified discriminative wavenumbers, indicated by dashed lines color-coded according to importance rank. (b) Normalized SHAP feature importance profile highlighting key Raman shifts contributing to the classification of the Pro:Val AA-AuNPs ratiometric mixtures.

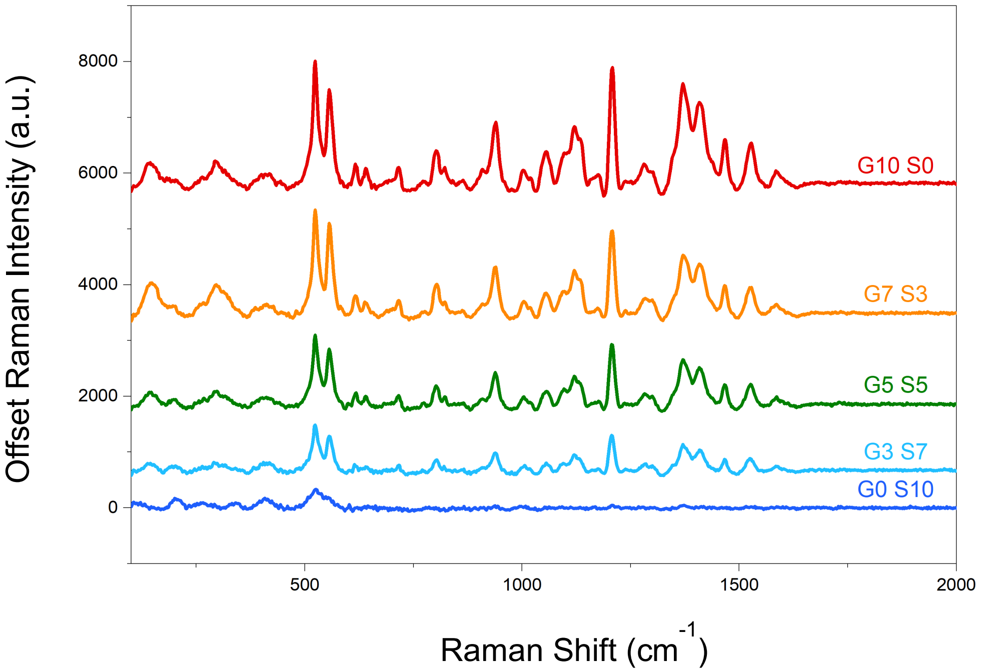


**Figure S18.** Representative SERS spectra of Glu:Ser AA-AuNPs ratiometric mixtures showing distinct spectra features at different Glu:Ser ratios.

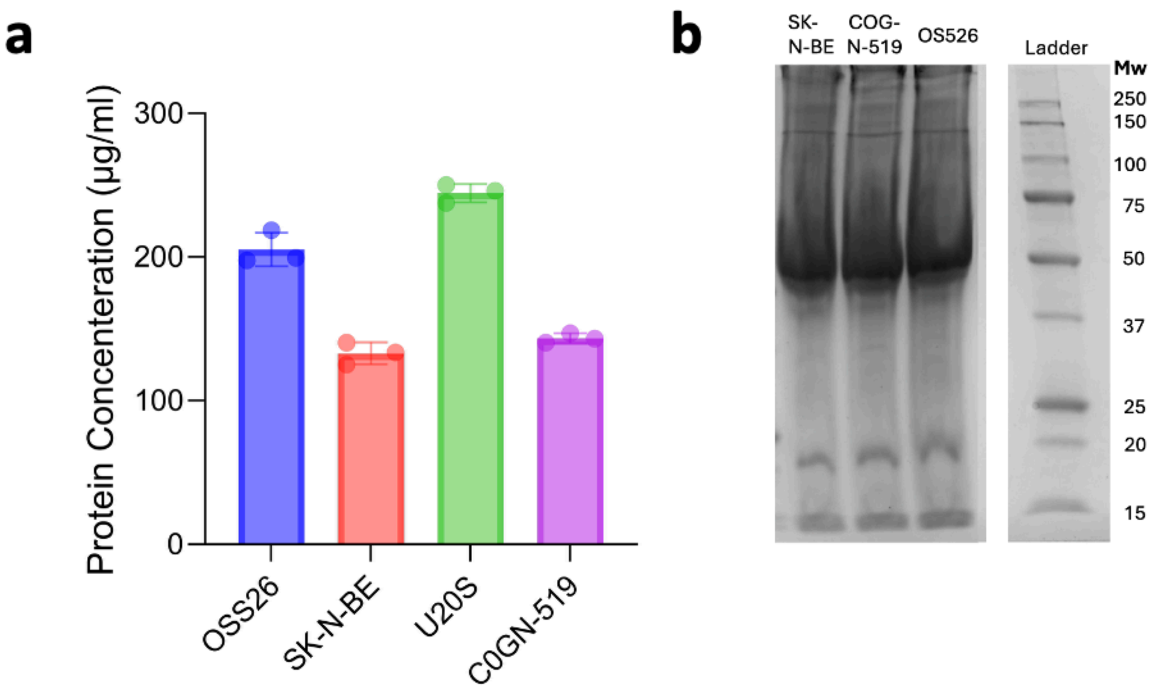


**Figure S19. Protein quantification and characterization of isolated exosomes.** (a) Protein concentration of exosomes isolated from four different cell lines. (b) SDS-PAGE analysis of proteins from isolated exosomes. Data are presented as mean ± standard deviation (n=2).

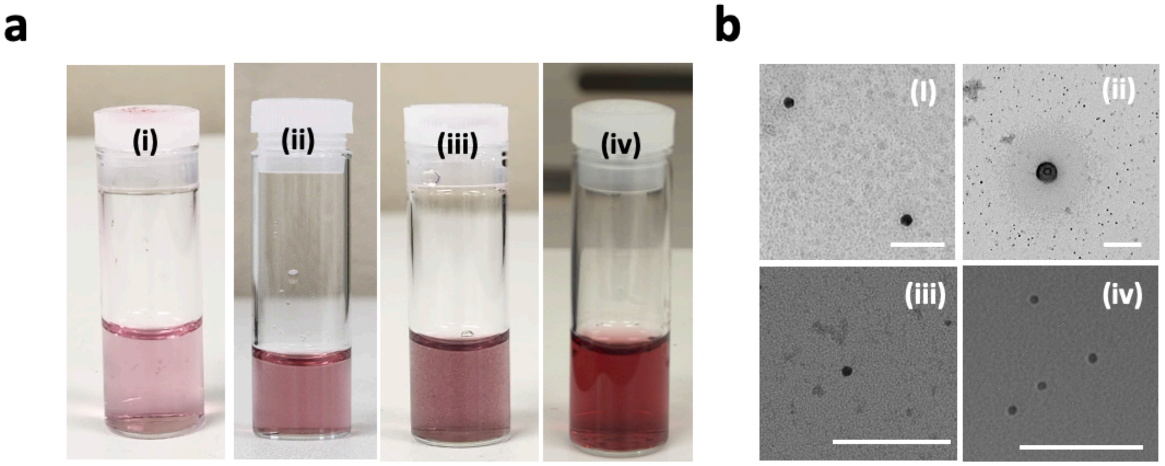


**Figure S20. Visual and morphological characterization of Exo-AuNPs.** (a) Digital photographs of Exo-AuNPs suspensions. (b) Representative negative-stained TEM images of (i) Exo-AuNP@OS-526, (ii) Exo-AuNP@COG-N-519, (iii) Exo-AuNP@SK-N-BE (2), and (iv) Exo-AuNP@U20 NPs. (scale bar: 100nm)

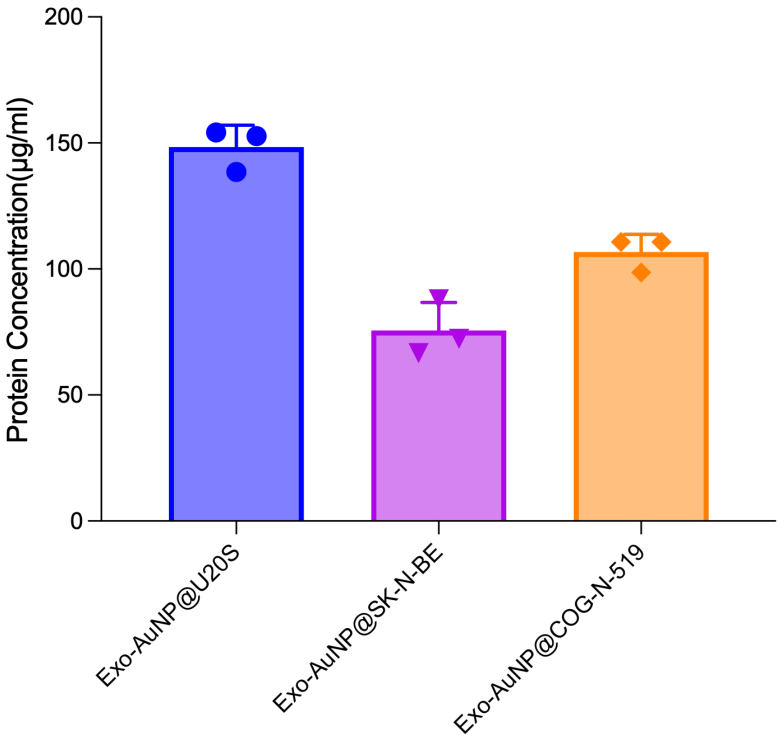


**Figure S21.** Protein concentration of the Exo-AuNPs determined *via* bicinchoninic acid (BCA) assay.

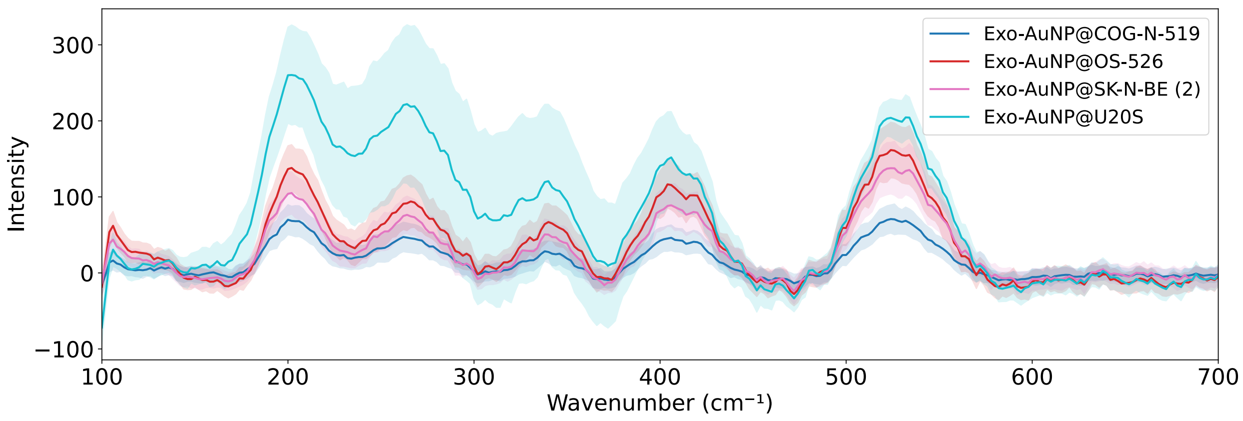


**Figure S22.** Mean SERS spectra (± standard deviation) for different Exo-AuNPs formulations.
